# Genome-wide CRISPRi/a screens in human neurons link lysosomal failure to ferroptosis

**DOI:** 10.1101/2020.06.27.175679

**Authors:** Ruilin Tian, Anthony Abarientos, Jason Hong, Sayed Hadi Hashemi, Rui Yan, Nina Dräger, Kun Leng, Mike A. Nalls, Andrew B. Singleton, Ke Xu, Faraz Faghri, Martin Kampmann

**Affiliations:** Institute for Neurodegenerative Diseases, Department of Biochemistry and Biophysics, University of California, San Francisco, San Francisco, CA, USA; Biophysics Graduate Program, University of California, San Francisco, San Francisco, CA, USA; Department of Computer Science, University of Illinois at Urbana-Champaign, Urbana, IL, USA; Department of Chemistry, University of California, Berkeley, Berkeley, CA, USA; Laboratory of Neurogenetics, National Institute on Aging, National Institutes of Health, Bethesda, MD, USA; Data Tecnica International, LLC, Glen Echo, MD, USA; Chan Zuckerberg Biohub, San Francisco, CA, USA

## Abstract

Single-cell transcriptomics provide a systematic map of gene expression in different human cell types. The next challenge is to systematically understand cell-type specific gene function. The integration of CRISPR-based functional genomics and stem cell technology enables the scalable interrogation of gene function in differentiated human cells. Here, we present the first genomewide CRISPR interference and CRISPR activation screens in human neurons.

We uncover pathways controlling neuronal response to chronic oxidative stress, which is implicated in neurodegenerative diseases. Unexpectedly, knockdown of the lysosomal protein prosaposin strongly sensitizes neurons, but not other cell types, to oxidative stress by triggering the formation of lipofuscin, a hallmark of aging, which traps iron, generating reactive oxygen species and triggering ferroptosis. We also determine transcriptomic changes in neurons following perturbation of genes linked to neurodegenerative diseases. To enable the systematic comparison of gene function across different human cell types, we establish a data commons named CRISPRbrain.

---

The human body comprises hundreds of different cell types. Even though their genomes are nearly identical, cell types are characterized by vastly different cell biologies, enabling them to fulfill diverse physiological functions. Transcriptomic profiling, fueled by recent advances in single-cell- and single-nucleus-RNA sequencing technologies, has enabled the establishment of a Human Cell Atlas of cell-type specific gene expression signatures^1^. In addition to gene expression, gene function can also be cell type-specific, as evidenced by the fact that mutations in broadly expressed or housekeeping genes can lead to cell-type specific defects and disease states. Striking examples are familial mutations causing neurodegenerative diseases, such as hereditary motor and sensory neuropathies (Charcot-Marie-Tooth disease), which can be caused by mutations in ubiquitously expressed trafficking factors (RAB7A, DNM2, DYNC1H1), or aminoacyl-tRNA synthetases (AARS, GARS, HARS, MARS)^2^.Moreover, neurodegenerative diseases are often characterized by the selective vulnerability of specific neuronal subtypes, even if the mutated gene is expressed throughout the brain or even throughout the whole body. Celltype specific gene function is also supported by our recent finding that knockdown of certain genes can have remarkably different impacts on cell survival and gene expression in different isogenic human cell types, including stem cells and neurons^3^.

Therefore, understanding the function of human genes in different cell types is the next step toward elucidating tissue-specific cell biology and uncovering disease mechanisms. To this end, we recently developed a functional genomics platform, leveraging the strengths of CRISPR interference (CRISPRi) and induced pluripotent stem cell (iPSC) technology, that enables large-scale, multimodal loss-of-function genetic screens in differentiated human cell types, as demonstrated in neurons^3^.

Here, we present a gain-of-function screening platform in human iPSC-derived neurons based on CRISPR activation (CRISPRa), which can yield complementary biological insights to CRISPRi screens^4^. Together, CRISPRi and CRISPRa in iPSC-derived neurons have enormous potential to uncover mechanisms and therapeutic targets in neurological diseases^5^. We conduct the first genome-wide CRISPRi and CRISPRa screens in human neurons based on a panel of cellular phenotypes, and CROP-seq screens to uncover transcriptional fingerprints of genes associated with neurodegenerative diseases. We apply our functional genomics platforms to systematically identify genetic modifiers of levels of reactive oxygen species and peroxidized lipids, and neuronal survival under oxidative stress, one of the predominant stresses in aging^6^ and neurodegenerative diseases^7^. These screens uncovered an unexpected role for the lysosomal protein prosaposin *(PSAP),* knockdown of which caused the formation of lipofuscin, a hallmark of aging, which traps iron, generating reactive oxygen species and triggering ferroptosis. Intriguingly, *PSAP* deficiency caused these dramatic phenotypes only in neurons, but not in iPSCs or HEK293 cells.

These results demonstrate the power of our approach in uncovering novel, cell-type specific human cell biology. We establish a Data Commons, named CRISPRbrain, for systematic exploration, interactive visualization, and comparison of functional genomics screening results in differentiated human cell types.

## Results

### Genome-wide CRISPRi and CRISPRa screens reveal genes controlling survival of human neurons

We previously established a CRISPRi platform that enables robust knockdown of endogenous genes and large-scale loss-of-function genetic screens in human iPSC-derived neurons^3^. Here, we expand our toolbox by establishing a CRISPRa platform for overexpression of endogenous genes and genome-wide gain-of-function screens in human iPSC-derived neurons. We adapted a previously published inducible CRISPRa system, DHFR-dCas9-VPH, whose function had been validated in human iPSCs^8^. In this system, the CRISPRa machinery is tagged by a DHFR degron leading to proteasomal degradation of the entire fusion protein. Trimethoprim (TMP) stabilizes the DHFR degron and prevents turnover, thereby inducing CRISPRa activity. As for our CRISPRi platform, an expression cassette for the CRISPRa machinery, CAG promoter-driven DHFR-dCas9-VPH, was stably integrated into the CLYBL safe-harbor locus of an iPSC line with an inducible Neurogenin 2 *(Ngn2)* expression cassette in the AAVS1 safe-harbor locus, which can be efficiently differentiated into homogenous glutamatergic neurons in a highly scalable manner^9^ (i^3^N-iPSC, Fig. 1a). A monoclonal line of these CRISPRa-iPSCs was generated and a normal karyotype was confirmed (Extended Data Fig. 1a). We validated the functionality of CRISPRa-iPSCs by confirming robust induction of an endogenous gene, *CXCR4,* in iPSC-derived neurons in a tightly inducible manner (Fig. 1b). CRISPRi and CRISPRa efficacies were comparable between iPSCs and derived neurons (Extended Data Fig. 1b,c).

**Fig. 1:**
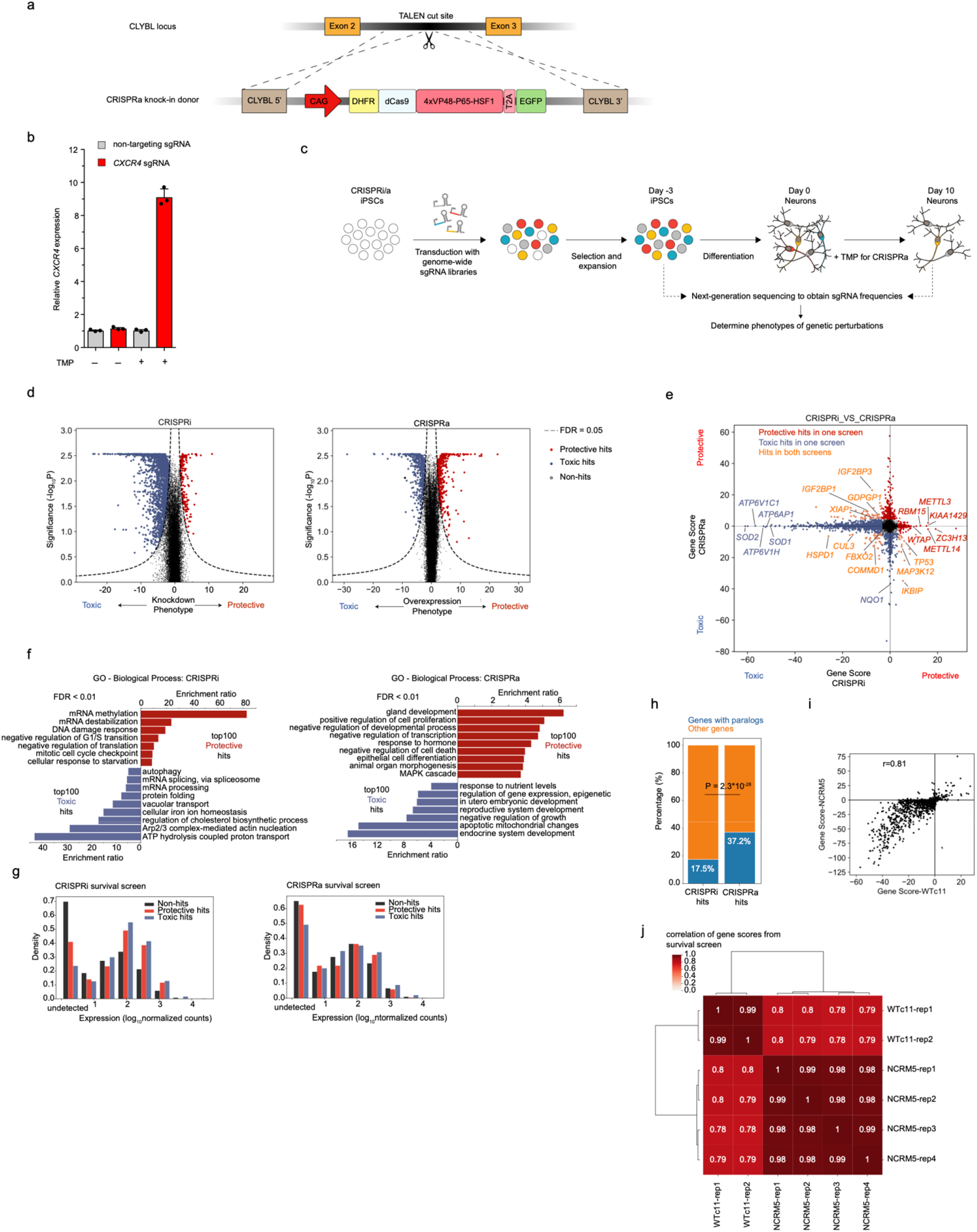
Genome-wide CRISPRi and CRISPRa screens in human iPSC-derived neurons identify regulators of neuronal survival. (a) Strategy for generating the CRISPRa iPSC line: an inducible CRISPRa construct, CAG promoter-driven DHFR-dCas9-VPH, was stably integrated into the CLYBL safe harbor locus through TALEN-mediated knock-in. dCas9, catalytically dead Cas9. VPH, activator domains containing 4X repeats of VP48, P65, and HSF1. (b) Functional validation of CRISPRa activity. qPCR quantification of the relative fold change of CXCR4 mRNA levels in CRISPRa-neurons expressing a CXCR4 sgRNA as compared to a nontargeting control sgRNA in the presence or absence of trimethoprim (TMP), which stabilizes the DHFR degron (mean +/− sd, n = 3 technical replicates). CXCR4 levels were normalized to the housekeeping gene ACTB. (c) Strategy for neuronal survival screens. CRISPRi/a iPSCs were transduced with genome-wide sgRNA libraries, containing ~100,000 sgRNAs targeting ~19,000 protein-coding genes and ~1,800 non-targeting control sgRNAs. TMP was added to CRISPRa neurons from Day 0 to induce CRISPRa activity. Frequencies of cells expressing a given sgRNA were determined by next-generation sequencing for Day 10 neurons and Day −3 iPSCs. (d) Volcano plots summarizing knockdown or overexpression phenotypes and statistical significance (Mann-Whitney U test) for genes targeted in the CRISPRi (left) and CRISPRa (right) screens. Dashed lines: False-discovery rate (FDR) cutoff for hit genes (FDR = 0.05) based on the Gene Score, see main text and Methods) (e) Comparing Gene Scores for hits from CRISPRi and CRISPRa screens. Hit genes with protective or toxic phenotypes in either screen are shown in red or blue, respectively. Genes that are hits in both screens are shown in orange. (f) Gene Ontology (GO) term enrichment analysis for the top 100 hit genes with protective or toxic phenotypes in CRISPRi (left) and CRISPRa (right) survival screens. Significantly enriched Biological Process terms (FDR < 0.01) are shown. (g) Expression levels of hit genes and non-hit genes from CRISPRi (left) or CRISPRa (right) screens are shown, binned by order of magnitude. (h) Percentage of hits with paralogs in CRISPRi and CRISPRa survival screens. A list of human paralog genes was obtained from a previous study^16^. P value was calculated using Fisher’s exact test. (i) Comparison of Gene Scores from survival screens in neurons derived from two different CRISPRi-iPSC lines, WTc11 (x-axis) and NCRM5 (y-axis), using a custom sgRNA library targeting 2,131 hit genes from the primary CRISPRi screen in WTc11. The Pearson correlation coefficient (r) is indicated. (j) Pairwise Pearson correlation of Gene Scores between biological replicates of survival screens in neurons derived from WTc11 and NCRM5 lines.

We previously conducted a sub-genome scale CRISPRi screen to reveal genes controlling neuronal survival using an sgRNA library targeting 2,325 genes representing the “druggable genome”^3^. Here, we greatly expanded the screen to target all protein-coding human genes in both loss-of- and gain-of-function screens by CRISPRi and CRISPRa, respectively. To our knowledge, these are the first genome-wide CRISPR screens in human neurons. We lentivirally transduced our CRISPRi and CRISPRa iPSCs with the genome-wide hCRISPRi/a-v2 sgRNA libraries^10^ and differentiated them into neurons (Fig. 1c). For the CRISPRa screen, TMP was added to Day 0 neurons to induce CRISPRa activity. Based on the depletion or enrichment of sgRNAs targeting specific genes at Day 10 compared to Day –3, we identified hit genes for which knockdown or overexpression was toxic or protective for neuronal survival (Fig. 1d, Supplementary Table 1), using our previously published pipeline, MAGeCK-iNC^3^. For each gene, a Gene Score was calculated to capture both the effect size of the phenotype and the statistical significance (see Methods). For most hit genes, only a subset of the 5 sgRNAs targeting the gene had a significant phenotype (Extended Data Fig. 1d, e), justifying the use of 5 sgRNAs per gene in the genomewide library. The likelihood of detecting hit genes was not related to the length of the gene or its coding sequence (Extended Data Fig. 1f,g).

For CRISPRi, among the top 10 hits with toxic knockdown phenotypes were genes encoding superoxide dismutases *SOD1* and *SOD2,* which protect cells from oxidative stress (Fig. 1e). Intriguingly, *SOD1* and *SOD2* are not broadly essential in other human cell types, including pluripotent stem cells^11–13^ and cancer cells^14^, suggesting a selective vulnerability to oxidative stress of neurons. Genes encoding subunits of the vacuolar ATPase (V-ATPase) complex, including *ATP6V1H, ATP6V1C1,* and *ATP6AP1,* were also among the top hits with toxic knockdown phenotypes. The V-ATPase complex mediates acidification of the endo-lysosomal compartment through ATP hydrolysis-coupled proton transport, and its dysfunction can cause neurodegenerative diseases^15^. Gene Ontology (GO) analysis revealed additional pathways that were enriched in the top 100 hits with toxic or protective phenotypes in the CRISPRi screen (Fig. 1f, left). For example, genes involved in cholesterol biosynthesis were strongly enriched in hits with toxic knockdown phenotypes, suggesting an important role of cholesterol in maintaining neuronal survival, consistent with our previous findings^3^. Other homeostatic pathways, including iron homeostasis, protein folding, mRNA processing, and autophagy, were also essential for neuronal survival.

For CRISPRa, GO analysis revealed that overexpression of pro- or anti-apoptotic genes showed expected toxic or protective survival phenotypes, respectively, validating our approach (Fig. 1f, right). Next, we compared hits from our CRISPRi and CRISPRa screens (Fig. 1e). Overall, there was little overlap between CRISPRi and CRISPRa hits, consistent with our previous study comparing parallel CRISPRi and CRISPRa screens in K562 cells^4^. The fact that CRISPRi and CRISPRa screens uncover distinct sets of hit genes can be explained by several factors. First, a gene that is not expressed in neurons will not have a CRISPRi knockdown phenotype, but may have a CRISPRa overexpression phenotype. Indeed, genes expressed at low or undetectable levels were strongly depleted from CRISPRi hits (Fig. 1g, left), whereas CRISPRa hits were not restricted by endogenous expression levels (Fig. 1g, right). Second, many genes encode proteins that form complexes (such as the V-ATPase complex), for which knockdown of a single component could abrogate the function of the entire complex and result in a phenotype, whereas overexpression of a single subunit by CRISPRa would generally be insufficient to induce an increased function of the complex. Last, knockdown of a single gene may not lead to a phenotype due to redundancy. For example, gene paralogs can compensate for each other’s loss-of-function. Paralogs tend to have less deleterious loss-of-function effect in CRISPR-Cas9 screens in different human cell lines^16^. Consistent with this idea, the percentage of hit genes with paralogs was much lower in our CRISPRi screen compared to that in CRISPRa screen (17.5% (355 / 2032) vs. 37.2% (314 / 845), P < 0.00001, Fig. 1h). Taken together, CRISPRi and CRISPRa screens can uncover complementary biological insights.

Nevertheless, we identified a number of overlapping hit genes in the two screens (Fig. 1e). Many of these genes showed opposing phenotypes on neuronal survival upon CRISPRa induction and CRISPRi repression. Among these genes, we identified known regulators of apoptosis, such as *TP53, IKBIP,* and factors specifically of neuronal survival, such as *XIAP^17–20^, GDPGP1^21^* and *MAP3K12’^3,22–24^*, as well as genes not previously implicated in neuronal survival, such as *IGF2BP1* and *IGF2BP3.*

For some genes, including several involved in protein homeostasis (e.g. *CUL3, FBXO2, COMMD1,* and *HSPD1),* perturbations in both directions were detrimental to neuronal survival (i.e. genes in the left lower quadrant of Fig. 1e), suggesting their endogenous expression levels are narrowly balanced for optimal survival.

While some CRISPRi and CRISPRa hits were also survival-relevant in other cell types based on previously reported screens, a substantial fraction of hits was neuron-specific (Extended Data Fig. 1h-j). Some CRISPRa hits were related to development of other cell types (Extended Data Fig. 1i), suggesting that ectopic expression of developmental factors affected cell type identity and thereby cell survival/proliferation, as previously observed in K562 cells^4^.

We confirmed the robustness of our screening platform by conducting focused CRISPRi screens using an sgRNA library targeting the top hits from the genome-wide screen in neurons derived from two distinct iPSC lines: WTc11 (used in the primary screen), and NCRM5. Phenotypes were highly correlated for neurons derived from these two lines (Fig. 1i). However, reproducibility between replicate screens using the same cell line was slightly higher than between screens in different lines (Fig. 1j), suggesting that some neuronal phenotypes depend on the genetic background.

### Genome-wide CRISPRi/a screens elucidate pathways controlling neuronal response to oxidative stress

Given the unique vulnerability of neurons to redox imbalance-induced oxidative stress, which is often found in the brain of patients with neurodegenerative diseases^25–29^, we sought to apply our functional genomics toolkit to systematically identify factors that are important for redox homeostasis and oxidative stress response in human neurons. We performed screens based on two strategies.

First, we conducted genome-wide CRISPRi and CRISPRa screens to identify modifiers of neuronal survival under oxidative stress conditions (Fig. 2a). Standard neuronal culture medium contains a combination of antioxidants, including vitamin E, vitamin E acetate, superoxide dismutase, catalase, and glutathione. To create an environment of chronic low-level oxidative stress, we cultured neurons in medium lacking the above antioxidants (–AO medium). We reasoned that compared to acute harsh treatments to induce reactive oxygen species (ROS), such as adding H2O2 or rotenone, –AO medium provided a more physiologically relevant approximation of chronic oxidative stress.

**Fig. 2:**
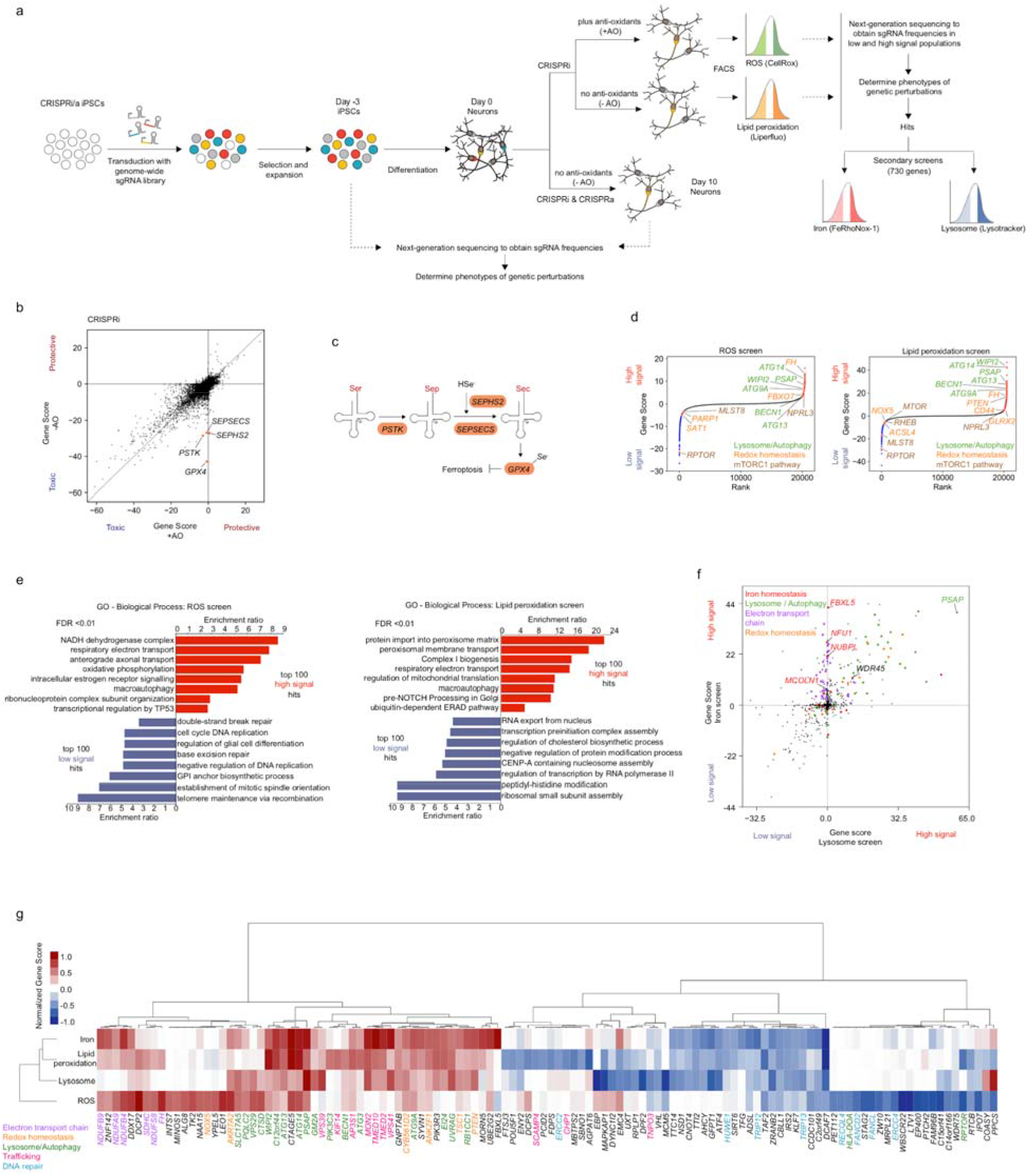
Genome-wide CRISPRi and CRISPRa screens in human iPSC-derived neurons identify regulators of oxidative stress survival and redox homeostasis. (a) Screening strategy. First, survival-based screens were conducted to identify modifiers of neuronal survival under mild oxidative stress induced by anti-oxidant (AO) removal from the neuronal medium (-AO). Second, FACS-based screens were conducted for modifiers of reactive oxygen species (ROS) and lipid peroxidation levels. Last, secondary screens for lysosomal status and labile iron levels were conducted to further characterize hit genes. (b) Comparison of Gene Scores in +AO and -AO conditions for ~19,000 protein-coding genes targeted in genome-wide CRISPRi survival screens. See main text and Methods for the definition of the Gene score. (c) Pathway for selenocysteine incorporation into GPX4. Hit genes are highlighted in orange. (d) Ranked Gene Scores from the ROS screen and the lipid peroxidation screen. High-signal hits are shown in red and low signal hits in blue. Genes discussed in the paper are highlighted in orange. (e) GO term enrichment analysis for the top 100 high-signal and low-signal hits in the ROS screen (left) and the lipid peroxidation screen (right). Significantly enriched Biological Process terms (false discovery rate (FDR) < 0.01) are shown. (f) Gene Scores from the lysosome and iron secondary screens targeting 730 genes selected from the primary genome-wide screens. Genes are color-coded by pathways based on Gene Ontology (GO) annotation. (g) Heatmap showing Gene Scores across screens (rows) for genes that are among the top 20 high-signal or low-signal hits in at least one screen (columns). Rows and columns are hierarchically clustered. Genes are color-coded by pathways based on GO annotation.

Next, we compared modifiers of neuronal survival in this oxidative stress condition to the modifiers of survival in the standard, unstressed condition (Fig. 2b, Supplementary Table 1). Interestingly, in the comparison of CRISPRi hits, we identified that *GPX4* (encoding the selenoprotein Glutathione Peroxidase 4) and genes responsible for selenocysteine incorporation into proteins (including *PSTK, SEPHS2,* and *SEPSECS)* were particularly essential for neurons to survive under oxidative stress (Fig. 2b,c). GPX4 utilizes glutathione to reduce peroxidized lipids and thus prevents ferroptosis, which is a non-apoptotic type of cell death caused by iron-dependent lipid peroxidation^30^. This result suggested that neurons could be susceptible to ferroptosis under oxidative stress conditions. Hits from CRISPRa screens showed high correlation (R = 0.82) between oxidative stress and unstressed conditions, suggesting no strong stress-specific phenotypes for overexpressed genes (Extended Data Fig. 2).

Second, we conducted genome-wide CRISPRi screens for modifiers of levels of ROS and peroxidized lipids in neurons. Specifically, we stained CRISPRi neurons transduced with genome-wide sgRNA libraries with fluorescent indicators of ROS and lipid peroxidation (CellRox and Liperfluo, respectively) and sorted them into high and low fluorescence populations by FACS (Fig. 2a). The MAGeCK-iNC pipeline was used to identify hit genes knockdown of which led to an increase (‘high signal”) or decrease (‘low signal’) in ROS or peroxidized lipids (Supplementary Table 1). From these screens, we identified both known and unexpected genetic modifiers of ROS and peroxidized lipid levels. We found that knockdown of components of the electron transport chain increased both ROS and lipid peroxidation levels (Fig. 2d). This was expected because ROS are mainly generated by proton leak from the electron transport chain, and knockdown of electron transport chain components could increase proton leakage. Many autophagy-related genes were also common hits in the two screens (Fig. 2 d,e), such as *WIPI2, ATG9A, ATG13, ATG14,* and *BECN1,* suggesting an important role of autophagy in maintaining redox homeostasis in cells, as reported in previous studies^31–33^.

Disruption of genes involved in the mTORC1 pathway, including components of the mTORC1 complex *(MTOR, RPTOR,* and *MLST8)* or its activator *RHEB* reduced ROS and/or lipid peroxidation levels, whereas knockdown of *NPRL3,* a mTORC1 inhibitor, induced ROS and lipid peroxidation levels in neurons (Fig. 2d). This is consistent with previous observations that increasing mTORC1 signaling induced ROS production^34^, whereas inhibiting mTORC1 reduced ROS^34–36^. Moreover, GO term enrichment analysis of lipid peroxidation hits revealed a strong enrichment of peroxisomal genes, knockdown of which increased lipid peroxidation, consistent with the important roles of peroxisomes in redox regulation and degradation of (poly-)unsaturated fatty acids.

*FBXO7,* a gene associated with Parkinson’s disease, whose deficiency was found to cause complex I respiratory impairment and ROS production^37^, also increased ROS levels when knocked down in our screen (Fig. 2d). Knockdown of other previously characterized ROS regulators, including positive regulators such as *PARP1^38^, SAT1^39^* and *NOX5^40^* and negative regulators such as *PTEN^41^* and *FH^42,43^* showed the expected effects on ROS and/or peroxidized lipid levels in our screens (Fig. 2d).

Interestingly, key regulators of ferroptosis were also hits in the lipid peroxidation screen. *ACSL4*, encoding Acyl-CoA Synthetase Long Chain Family Member 4, which enriches cellular membranes with long PUFAs is required for ferroptosis^44,45^. *ACLS4* inhibition has been shown to prevent ferroptosis^44^, consistent with a reduction of peroxidized lipids upon knockdown in our screen (Fig. 2d). By contrast, knockdown of *CD44,* whose splicing variant CD44v stabilizes the cystine/glutamate antiporter xCT at the plasma membrane and increases cysteine uptake for GSH synthesis, thereby inhibiting ferroptosis^46,47^, increased lipid peroxidation in our screen as expected (Fig. 2d).

Surprisingly, we found several strong hit genes with lysosomal functions but not associated with autophagy (Fig. 1d). In particular, *PSAP* (encoding the lysosomal protein prosaposin) was one of the strongest hits in both ROS and lipid peroxidation screens (Fig. 2d).

To further investigate hit genes from the genome-wide ROS and lipid peroxidation screens in high-throughput, we conducted focused secondary screens with an sgRNA library targeting 730 hit genes (Fig. 2a). One secondary screen used lysotracker stain as a readout to uncover redox hits that may also alter lysosomal status. Lysotracker is a fluorescent acidotropic probe for labeling and tracking acidic organelles in live cells, which has been used to detect lysosome localization, quantify the number and sizes of lysosomes^48,49^ and to monitor autophagy^50,51^. The other secondary screen used the labile Fe^2+^-detecting probe FeRhoNox-1. Since intracellular labile ferrous iron (Fe^2+^) can contribute to ROS generation and lipid peroxidation via Fenton reactions^52^, we asked whether some of the redox hits act through changing iron homeostasis.

Our secondary screens uncovered that several of our original ROS/lipid peroxidation hit genes also strongly affected iron and/or lysosome status (Fig. 2f, Supplementary Tables 1, 2). We found that knockdown of many lysosome/autophagy-related genes affected both lysosomal status and iron levels, reflecting the key role of lysosomes in iron homeostasis^53,54^. Among these genes, we found *WDR45,* which is involved in autophagosome formation and lysosomal degradation and is also associated with Neurodegeneration with Brain Iron Accumulation (NBIA) in line with previous studies^55,56^. We also identified other known iron regulators for which knockdown increased iron levels, including *FBXL5,* a negative regulator of iron levels^57^, the Fe/S cluster biogenesis genes *NFU1* and *NUBPL,* and *MCOLN1,* which encodes an endolysosomal iron release channel^58^ (Fig. 2f). Interestingly, knockdown of genes involved in the electron transport chain increased labile iron (Fig. 2f). Mitochondria are a major site of cellular iron storage and utilization in processes such as heme synthesis and iron-sulfur cluster biogenesis^59^. Disruption of the electron transport chain may lead to mitochondrial damage which can cause the subsequent release of labile iron and reduced iron utilization, thus increasing labile iron levels.

In summary, our screens for survival of oxidative stress and levels of ROS and peroxidized lipids uncovered many categories of known redox regulators, validating the sensitivity of our approach and supporting the notion that core mechanisms of redox regulation are conserved across different cell types. A substantial fraction of the hit genes were also modifiers of labile iron levels and/or lysosomal status (Fig. 2g).

### Loss of prosaposin increases ROS and lipid peroxidation levels in neurons and causes neuronal ferroptosis under oxidative stress

Surprisingly, we identified that knockdown of *PSAP* altered lysosomal status and strongly induced ROS, lipid peroxidation, and iron levels when depleted (Fig. 2d,f,g). Given the screen phenotypes of *PSAP,* which were unexpected based on its known functions, and the association of *PSAP* variants not only with lysosomal storage disorders^60–62^ but also Parkinson’s Disease^63^, we decided to further investigate the underlying mechanisms of *PSAP* in neuronal redox regulation.

*PSAP* encodes prosaposin, which is a pro-protein that is proteolytically processed by cathepsin D (encoded by *CTSD*) in the lysosome to generate four cleavage products: saposins A, B, C, and D^64^. These four saposins, along with the lysosomal protein GM2A (GM2 Ganglioside Activator), function as activators for glycosphingolipid degradation by lysosomal hydrolases^65^ (Fig. 3a). Intriguingly, both *CTSD* and *GM2A* were also hits in at least one of the redox screens, showing similar knockdown phenotypes as *PSAP* (Fig. 3b). Together, these findings suggest an important and unexpected role for glycosphingolipid degradation in redox homeostasis.

**Fig. 3:**
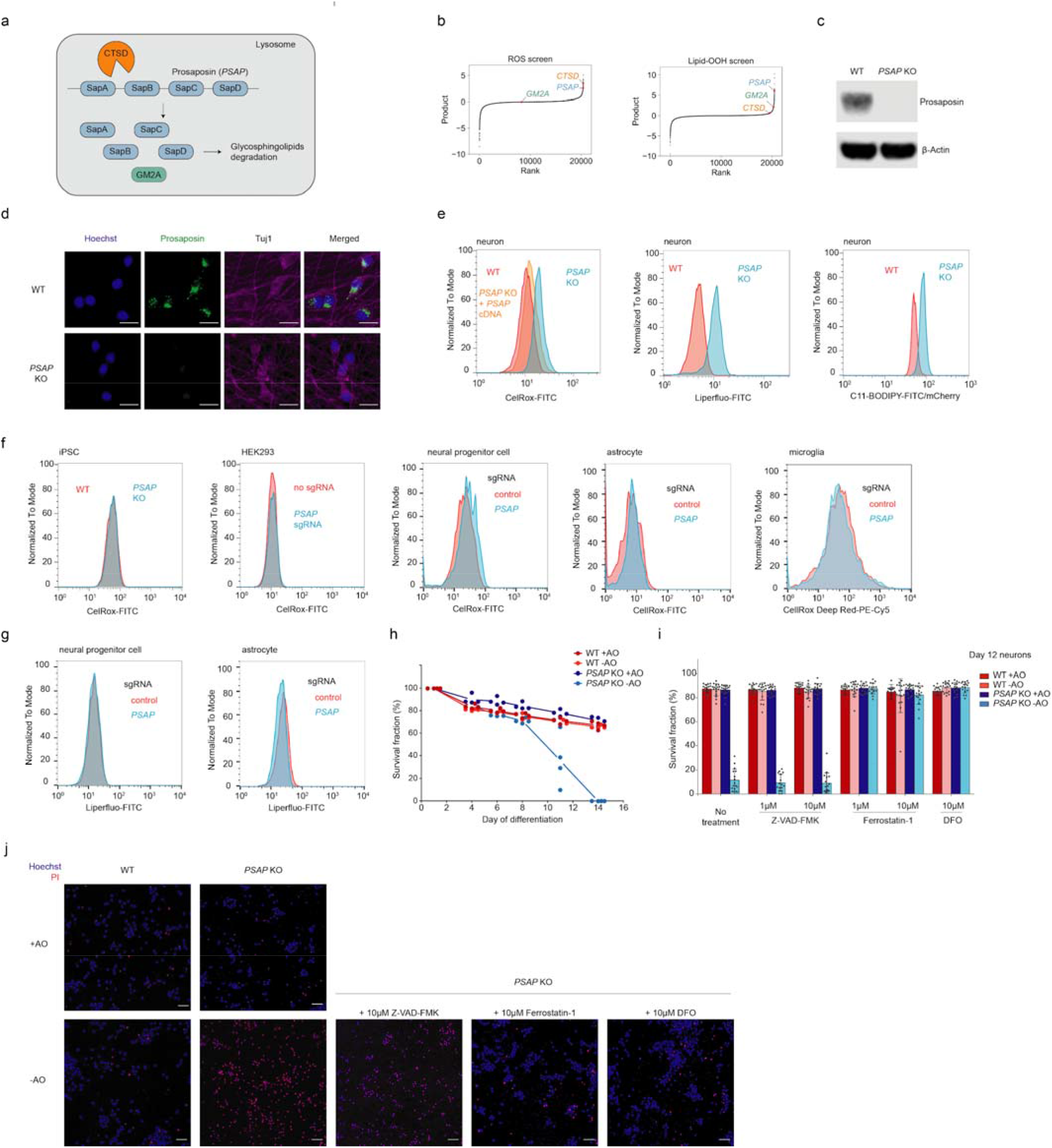
Loss of prosaposin induces ROS and lipid peroxidation in neurons and causes neuronal ferroptosis in the absence of antioxidants. (a) Prosaposin is processed in the lysosome by cathepsin D (encoded by *CTSD)* into saposin subunits, which function together with GM2A as activators for glycosphingolipid degradation. (b) Results from the reactive oxygen species (ROS) and lipid peroxidation screens (Fig. 2), highlighting *PSAP* and the related genes *CTSD* and *GM2A.* (c) Western blot showing the depletion of prosaposin in the *PSAP* knockout (KO) iPSC line. (d) Representative immunofluorescence microscopy images showing the loss of prosaposin in *PSAP* KO neurons. WT and *PSAP* KO neurons were fixed and stained by antibodies against prosaposin (shown in green) and the neuronal marker Tuj1 (shown in purple). Nuclei were counterstained by Hoechst, shown in blue. Scale bars, 20 μm. (e) ROS levels (as indicated by CellRox) and lipid peroxidation levels (as indicated by Liperfluo and C11-BODIPY) in WT and *PSAP* KO neurons, measured by flow cytometry. (f) ROS levels in iPSCs, HEK293 cells, neural progenitor cells, astrocytes and microglia in WT and *PSAP* KO backgrounds *(PSAP* KO for iPSCs and *PSAP* knockdown by CRISPRi in the other cell types), measured by CellRox via flow cytometry. (g) Lipid peroxidation levels (as indicated by Liperfluo) in WT and PSAP knockdown neural progenitor cells and astrocytes, measured by flow cytometry. (h) Survival curves for WT and *PSAP* KO neurons cultured in normal neuronal medium (+AO) or medium lack of antioxidants (-AO), quantified by imaging using Hoechst stain (all cells) and propidium iodide (PI) (dead cells). Survival fraction is calculated as (total cell count – dead cell count) / total cell count. Data is shown as mean +/− sd, n = 4 culture wells per group. 16 imaging fields were averaged for each well. (i) Survival fractions of WT and *PSAP* KO neurons treated with different cell death inhibitors under +AO or - AO conditions, quantified by imaging in the same way as for G. Data is shown as mean +/− sd, n = 16 imaging fields per group. (j) Representative images for the Hoechst (shown in blue) and propidium iodide (PI, shown in red) staining in h. Scale bars, 50 μm.

To validate our screen results using an independent approach, we generated a clonal *PSAP* knockout (KO) iPSC line (Fig. 3c,d). Next, we measured ROS and lipid peroxidation levels in wild-type (WT) and *PSAP* knockout (KO) neurons. We included C11-BODIPY, another indicator for peroxidized lipid, as additional validation for lipid peroxidation levels. Indeed, we observed a substantial increase in ROS and peroxidized lipid levels in *PSAP* KO neurons, confirming our screen results (Fig. 3e). Moreover, ROS induction in *PSAP* KO neurons could be rescued by the overexpression of *PSAP* cDNA (Fig. 3e), confirming that phenotypes were not driven by off-target genome editing. Interestingly, *PSAP* KO or knockdown in five other cell types we tested (HEK293 cells, iPSCs, and iPSC-derived neuronal progenitor cells, astrocytes and microglia, Extended Data Fig. 3a) did not increase ROS levels (Fig. 3f), suggesting a neuron-specific role for *PSAP* in redox regulation. Similarly *PSAP* knockdown did not increase peroxidized lipid levels in two cell types we tested (Fig. 3g).

Next, we asked if increased ROS in *PSAP* KO neurons affects survival. We did not observe a survival defect of *PSAP* KO neurons over two weeks of culture in standard neuronal medium (Fig. 3h). Strikingly, however, when we cultured these neurons in medium lacking antioxidants (–AO), *PSAP* KO caused a dramatic decrease in survival around Day 11, resulting in complete death of all neurons by Day 14 (Fig. 3h).

To further investigate the underlying mechanism of cell death, we treated WT and *PSAP* KO neurons with compounds that inhibit different cell death pathways. Intriguingly, the viability of *PSAP* KO neurons under the -AO condition was not rescued by Z-VAD-FMK, a pan-caspase inhibitor that blocks apoptosis, but was fully rescued by ferroptosis inhibitors, including the iron chelator deferoxamine (DFO) and the lipid peroxidation inhibitor ferrostatin-1 (Fig. 3i,j), suggesting that mild oxidative stress triggers ferroptosis in *PSAP* KO neurons.

In summary, we validated the unexpected strong hit gene from our unbiased screens, *PSAP*, as a strong redox modifier in human neurons. Loss of *PSAP* increases ROS levels and leads to increased lipid peroxidation, resulting in neuronal ferroptosis in response to mild oxidative stress.

### Loss of prosaposin leads to lipofuscin formation and iron accumulation

Given the surprising connection between *PSAP* and ferroptosis, we further investigated the underlying mechanism. Neurons in the following experiments were cultured in standard neuronal medium (+AO) unless otherwise specified. We first asked if the loss of *PSAP* blocked glycosphingolipid degradation, given the canonical function of saposins. Indeed, untargeted lipidomics revealed that almost all glycosphingolipid species accumulated significantly in *PSAP* KO neurons compared to WT neurons (Supplementary Table 3, FDR < 0.01, Fig. 4a,b). Ether lipids, which are peroxisome-derived glycerophospholipids, were also enriched in *PSAP* KO neurons. The mechanisms driving ether lipid accumulation and whether this accumulation is protective or maladaptive remain to be investigated. Interestingly, the accumulation of ether lipids was also recently characterized as a feature of hypoxia^66^.

**Fig. 4:**
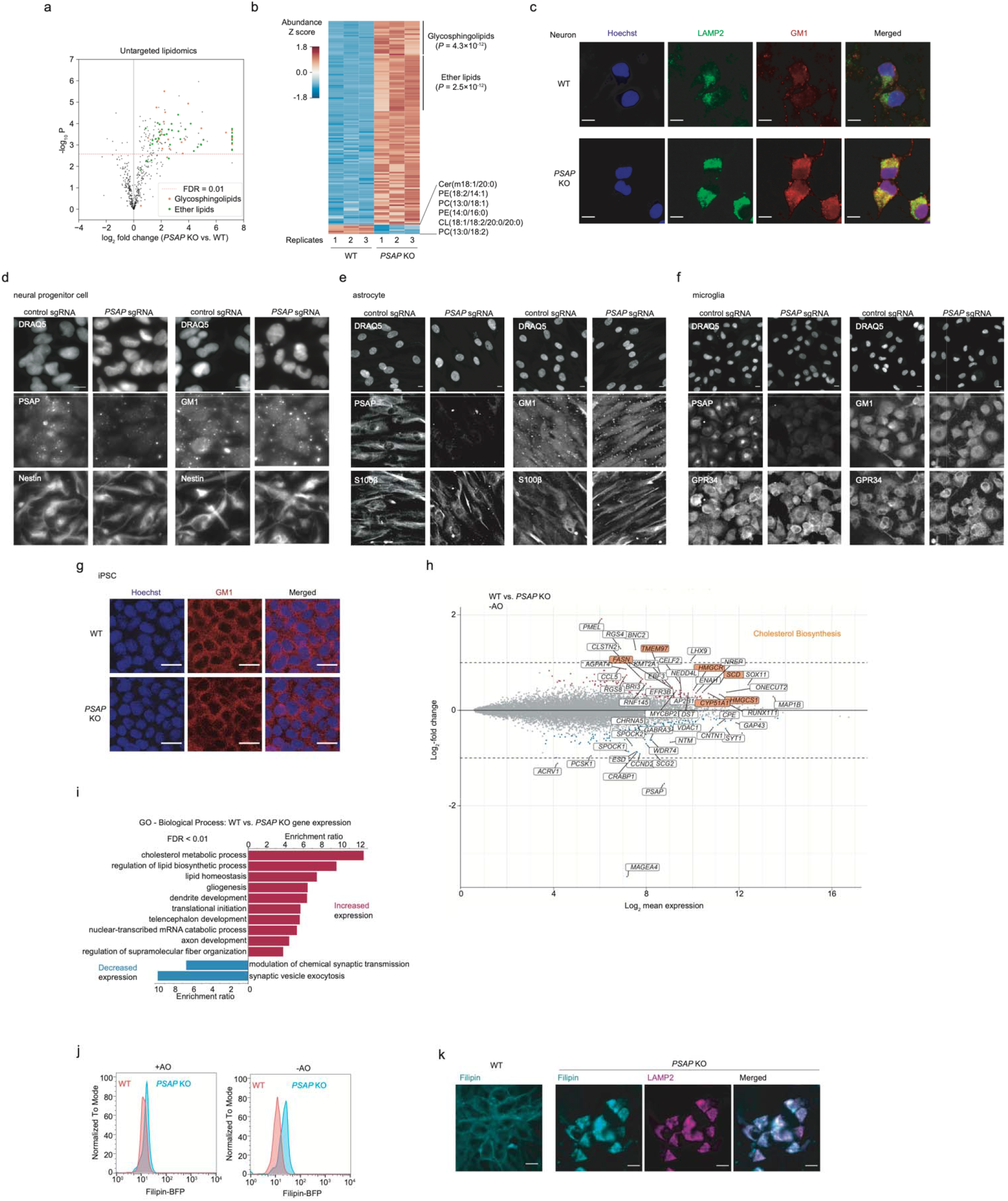
Loss of prosaposin disrupts glycosphingolipid degradation specifically in neurons but not other cell types, and leads to cholesterol accumulation. (a) Untargeted lipidomics comparing abundances of different lipid species in wild-type (WT) and PSAP knockout (KO) neurons. P values were calculated using two-sided Student’s t-test (n = 3 replicates per group). Dashed line, P value cutoff for false-discovery rate (FDR) < 0.01. Glycosphingolipids are shown in orange and ether lipids in green. (b) Heatmap showing the abundances of significantly increased or decreased lipids in PSAP KO neurons as compared to WT (FDR < 0.01). Enrichment P values for glycosphingolipids and ether lipids were calculated using Fisher’s exact test. Lipid abundances were Z score-normalized across samples. (c) Representative immunofluorescence microscopy images for WT and PSAP KO neurons stained with LAMP2 antibodies (shown in green) and GM1 antibodies (shown in red). Nuclei were counterstained by Hoechst, shown in blue. Scale bar, 10 μm. (d-f) Representative fluorescence microscopy images for neural progenitor cells (d), astrocytes (e) and microglia (f) diffentiated from CRISPRi iPSCs expression a non-targeting sgRNA or a PSAP sgRNA, stained with PSAP (left) and GM1 (right) antibodies. DRAQ5 was used for nuclear staining. Nestin, S100β and GPR34 antibodies were used as markers for neural progenitor cells, astrocytes and microglia, respectively. Scale bar, 10 μm. (g) Representative immunofluorescence microscopy images for WT and PSAP KO iPSCs stained with GM1 antibodies (shown in red). Nuclei were counterstained by Hoechst, shown in blue. Scale bar, 20 μm (h) Gene expression changes in PSAP KO neurons as compared to WT. Genes that are significantly upregulated and downregulated in PSAP KO neurons are shown in red and blue, respectively (false-discovery rate (FDR) < 0.05). The top 50 up- and down-regulated genes are labeled, and within this set, genes involved in the cholesterol biosynthesis pathway are highlighted in orange. (i) Gene ontology (GO) term enrichment analysis for significantly up- and down-regulated genes (FDR < 0.05) in PSAP KO neurons. Significantly enriched Biological Process terms are shown (FDR < 0.01). (j) Cholesterol levels measured by flow cytometry in filipin-stained WT and PSAP KO neurons cultured in the presence or absence of antioxidants (in +AO and -AO conditions). (k) Representative fluorescence microscopy images of WT and PSAP KO neurons stained with filipin (shown in cyan) and LAMP2 antibodies (shown for PSAP KO neurons, in red). Scale bar, 10 μm.

We also confirmed the accumulation of a specific glycosphingolipid species, GM1 ganglioside, in *PSAP* KO neurons by immunostaining (Fig. 4c). Interestingly, we did not observe GM1 accumulation in *PSAP* KO or knockdown iPSCs or iPSC-derived neuronal progenitor cells, astrocytes or microglia (Fig. 4d-g), suggesting a cell-type specific role of *PSAP.* Moreover, we performed RNA sequencing on WT and *PSAP* KO neurons. Interestingly, a number of genes involved in cholesterol biosynthesis were upregulated in *PSAP* KO neurons (Fig. 4h,i). Consistent with the induction of this pathway, we found increased cholesterol levels in lysosomes of *PSAP* KO neurons, as measured by Filipin staining (Fig. 4j,k). This finding mirrored the results of previous studies that *PSAP* is a strong genetic modifier of cholesterol levels^67^ and that accumulated glycosphingolipids can lead to the accumulation of cholesterol in lysosomes by inhibiting cholesterol efflux^68,69^.

Strikingly, we observed dramatically enlarged lysosomes in *PSAP* KO neurons (Fig. 4c), which were also reflected in an increased lysotracker signal by flow cytometry (Fig. 5a), as expected based on our screen results (Fig. 2f). Again, these lysosomal phenotype were not observed in HEK293 cells, iPSCs and derived astrocytes and microglia, and only to a small extent in iPSC-derived neuronal progenitor cells (Fig. 5b).

**Fig. 5:**
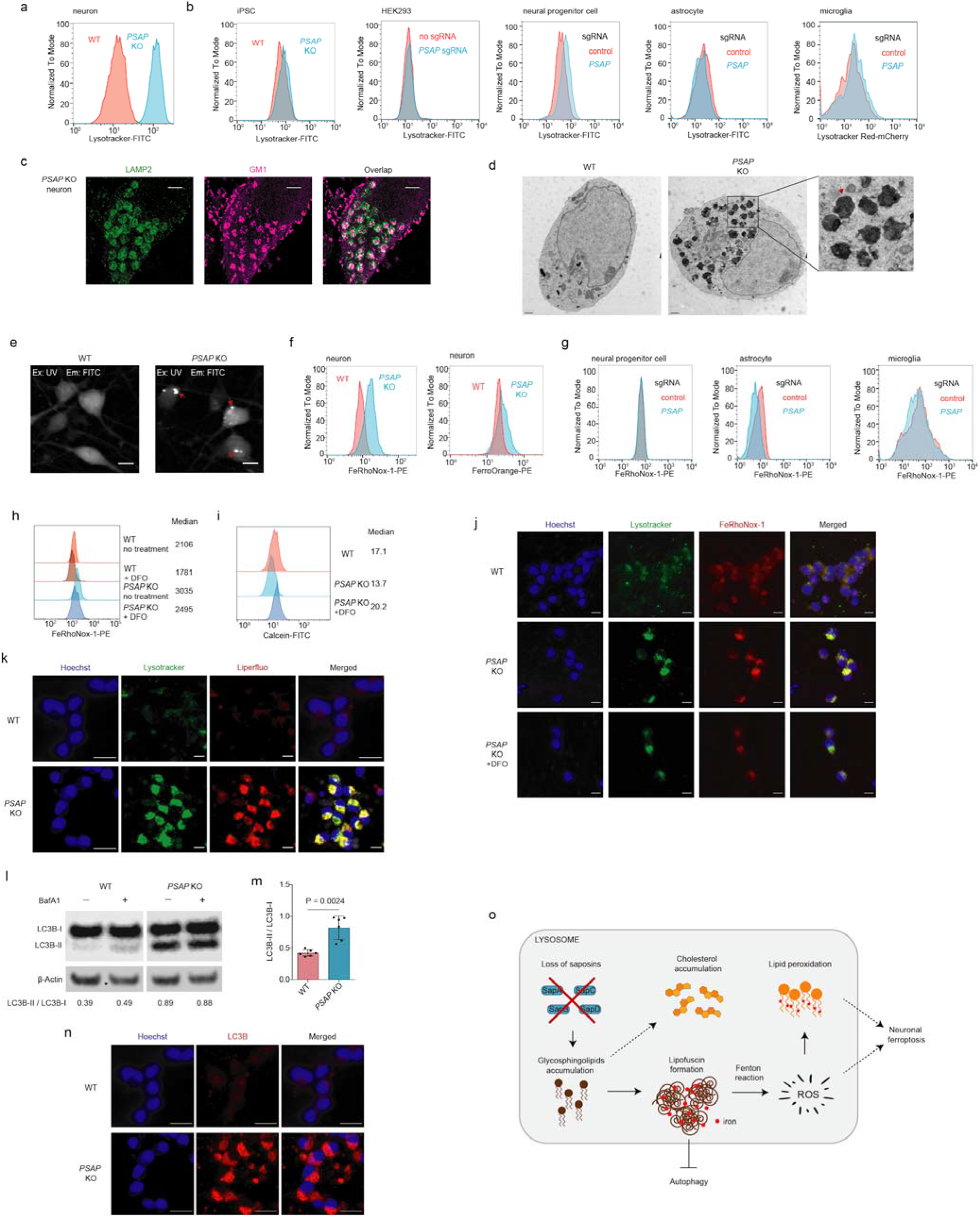
Impaired lipid degradation in PSAP KO neurons leads to lipofuscin formation, iron accumulation and impaired autophagy. (a) Lysotracker signals measured by flow cytometry in WT and *PSAP* KO neurons (b) Lysotracker signals measured by flow cytometry in iPSCs, HEK293 cells, neural progenitor cells, astrocytes and microglia (left to right) in WT and *PSAP* KO or knockdown backgrounds (*PSAP* KO for iPSCs and *PSAP* knockdown by CRISPRi in the other cell types) (c) Two-color STORM super-resolution images of PSAP KO neurons immunolabeled for LAMP2 (shown in green) and GM1 (shown in magenta). Scale bars, 2 μm. (d) Electron microscopy images for WT and PSAP KO neurons. The red arrow points to a representative lipofuscin structure. Scale bar, 1 μm. (e) Representative images for autofluorescence in WT and PSAP KO neurons. Excitation, UV (405 nm). Emission, FITC (525/20 nm). Scale bar, 10 μm. (f) Labile iron levels in WT and *PSAP* KO neurons. Neurons were stained with the iron indicators FeRhoNox-1 (left) or FerroOrange (right) and fluorescence was measured by flow cytometry. (g) Labile iron levels (as indicated by FeRhoNox-1) in WT and *PSAP* knockdown neural progenitor cells, astrocytes and microglia (left to right), measured by flow cytometry. (h) Labile iron levels in WT and *PSAP* KO neurons with or without DFO treatment. Cells were stained by FeRhoNox-1 and fluorescence was quantified by flow cytometry. Median signal intensities are indicated. (i) Labile iron levels in WT neurons, *PSAP* KO neurons, and *PSAP* KO neurons with DFO treatment. Cells were stained by Calcein and fluorescence was quantified by flow cytometry. Median signal intensities are indicated. (j) Representative fluorescence microscopy images for WT neurons, *PSAP* KO neurons, and *PSAP* KO neurons treated with 10 μM DFO for 3 days, stained with Lysotracker (shown in green) and FeRhoNox-1 (shown in red). Nuclei were counterstained by Hoechst, shown in blue. Scale bar, 10 μm. (k) Representative fluorescence microscopy images for WT and *PSAP* KO neurons under the - AO condition stained with Lysotracker (shown in green) and Liperfluo (shown in red). Nuclei were counterstained by Hoechst, shown in blue. Scale bar, 10 μm. (l) Western blot showing protein levels of phosphatidylethanolamine (PE)-conjugated LC3B (LC3B-II) and unconjugated LC3B (LC3B-I) in WT and *PSAP* KO neurons in the absence or presence of Bafilomycin A1 (BafA1). ß-Actin was used as a loading control. Ratios of LC3B-II to LC3B-I are indicated at the bottom. (m) Quantification of LC3B-II / LC3B-I ratios for WT and *PSAP* KO neurons (mean +/− sd, n = 6 independent experiments). P value was calculated using Student’s t-test. (n) Representative immunofluorescence microscopy images for WT and *PSAP* KO neurons, stained with LC3B antibodies (shown in red). Nuclei were counterstained by Hoechst, shown in blue. Scale bar, 20 μm. (o) A model for the mechanism linking prosaposin loss to neuronal ferroptosis. Loss of saposins blocks glycosphingolipid degradation in the lysosome. The build-up of glycosphingolipids leads to lipofuscin formation, which accumulates iron and generates reactive oxygen species (ROS) through the Fenton reaction. ROS then peroxidize lipids and cause neuronal ferroptosis in the absence of antioxidants. Other consequences are cholesterol accumulation and impaired autophagy.

To further characterize the enlarged lysosome-like structures accumulating in *PSAP* KO neurons, we performed STORM super-resolution microscopy^70^ and electron microscopy (EM). Similarly to conventional confocal microscopy (Fig. 4c), super-resolution microscopy revealed colocalization of accumulated GM1 and lysosomes (Fig. 5c), consistent with the notion that the lysosome is the main site of glycosphingolipid degradation. By EM, we remarkably observed a large number of electron-dense granules (Fig. 5d) that resembled the structure of lipofuscin (also known as age pigment). Lipofuscin is an insoluble aggregate of peroxidized lipids, proteins and metals in lysosomes of postmitotic cells, such as neurons, which is a hallmark of aging and also associated with several neurodegenerative diseases^71^. There is currently no consensus on the molecular mechanisms driving lipofuscin formation^71^. Further supporting the formation of lipofuscin in *PSAP* KO neurons was the detection of strong autofluorescence (Fig. 5e), a hallmark of lipofuscin^72^. Lipofuscin is known to accumulate iron from lysosomal degradation of iron-rich proteins or organelles (e.g. mitochondria), and accumulated iron in lipofuscin has been proposed to generate ROS through the Fenton reaction^73,74^. Indeed, we observed that *PSAP* deficiency caused a substantial increase of iron levels in neurons (Fig. 5f), but not other cell types (Fig. 5g). This result was consistent with our finding of *PSAP* as a top hit in our iron level screen (Fig. 2f). Iron accumulation occurred in lysosomes and was partially reversed by the iron chelator DFO (Fig. 5h-j). Meanwhile, dramatically increased lipid peroxidation was detected in lysosomes of *PSAP* KO neurons under the -AO condition (Fig. 5k), likely due to the accumulated iron, consistent with the strong lipid peroxidation-inducing phenotype of *PSAP* knockdown in our screen (Fig. 2d).

Other lysosomal functions were also disrupted in *PSAP* KO neurons, likely due to the formation of lipofuscin. We observed a massive accumulation of autophagosomes in *PSAP* KO neurons as indicated by the increased ratio of PE-conjugated form of LC3B to its unconjugated form (LC3B-II / LC3B-I) by western blot (Fig. 5l,m), and increased LC3B puncta by immunostaining (Fig. 5n). This substantial induction of LC3B puncta was not observed in other cell types upon *PSAP* knockdown (Extended Data Fig. 3b). Treatment with Bafilomycin A1 (BafA1), an inhibitor for degradation of autophagosomes by lysosomes, increased the LC3B-II / LC3B-I ratio in WT, but not *PSAP* KO neurons (Fig. 5l), suggesting a blockade of autophagic flux in *PSAP KO* neurons.

In summary, we elucidated that loss of saposins in *PSAP* KO neurons blocks glycosphingolipid degradation in lysosomes, leading to the formation of lipofuscin, which in turn accumulates iron and generates ROS that peroxidized lipids. The accumulation of peroxidized lipids leads to neuronal ferroptosis in the absence of antioxidants (Fig. 5o).

### Transcriptomic signatures of perturbations of disease-associated genes in human neurons

Over the past decade, genome-wide association studies (GWASs) have uncovered hundreds of genes that are associated with human neurodegenerative diseases^75,76^. However, functional characterizations of these risk genes are largely lacking^77^. Our CRISPRi and CRISPRa platforms provide a high-throughput approach to systematically interrogate gene function in human neurons. Beyond one-dimensional phenotypes such as survival or fluorescent reporter levels, CRISPR perturbation can be coupled to single-cell RNA sequencing, using CROP-seq or Perturb-Seq strategies^78–80^, to provide rich transcriptomic phenotypes.

The hit genes from our unbiased genome-wide CRISPRi and CRISPRa screens included hundreds of genes associated with neurodegenerative diseases (based on DisGeNet annotation^81^ and literature research). To better characterize these neurodegenerative disease risk genes, we performed CROP-seq experiments, targeting 184 genes for CRISPRi and 100 genes for CRISPRa with 2 sgRNAs per gene (Fig. 6a, see Method for details). Significant on-target gene knockdown or overexpression were detected for most of the targets in the CRISPRi or CRISPRa library, respectively (> 70% at P < 0.05, Fig. 6b). Within the population of cells expressing sgRNAs for a specific target gene, the levels of target knockdown or overexpression were heterogeneous in some cases (Fig. 6c,d), and we established a machine-learning based strategy to identify cells in which the targeted gene was effectively knocked down or overexpressed (Fig. 6c,d, Extended Data Fig 4a,b, see Methods for details).

**Fig. 6:**
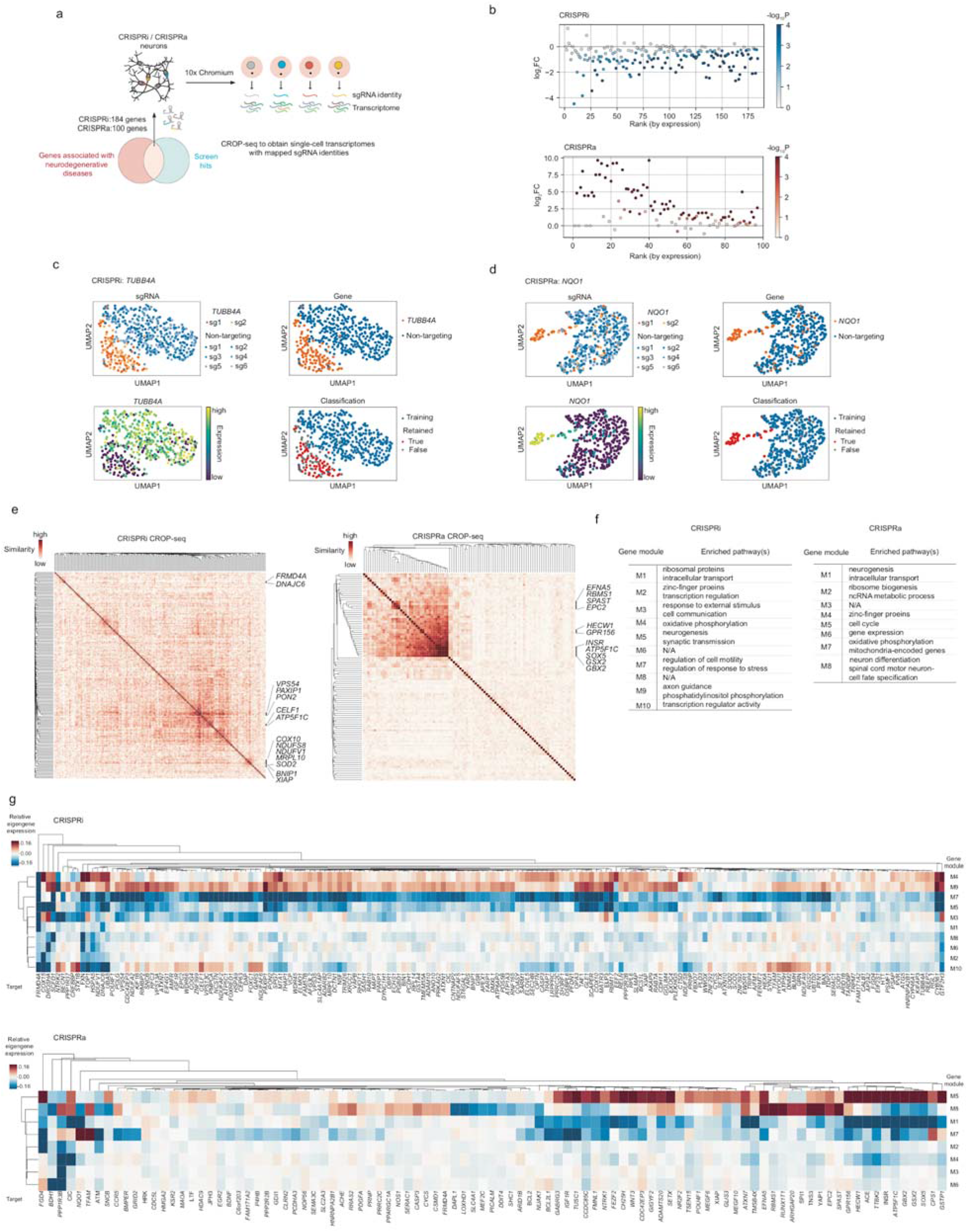
CROP-seq reveals transcriptomic responses to perturbations of neurodegenerative disease-associated genes in human iPSC-derived neurons. (a) Hit genes from our screens that are also associated with neurodegenerative diseases were targeted in a CROP-seq screen to detect their knockdown or overexpression effects on gene expression at single-cell resolution in human iPSC-derived neurons. (b) Summary of on-target knockdown by CRISPRi (top) or overexpression by CRISPRa (bottom) for all target genes in the CROP-seq libraries. log2FC represents the log2-fold change of the mean expression of a target gene in perturbed cells (i.e. cells expressing sgRNAs targeting that gene) compared to unperturbed cells (i.e. cells expressing non-targeting control sgRNAs). P values were calculated by the two-sided Wilcoxon rank-sum test. Target genes are ranked by their expression in unperturbed cells. (c,d) Examples of CROP-seq results showing on-target knockdown *(TUBB4A* in CRISPRi, c) or overexpression *(NQO1* in CRISPRa, d) and the classification method, shown in a twodimensional UMAP projection. (e) Pairwise similarities of differentially expressed genes among perturbations. Similarity scores were determined by the OrderedList package in R (see Methods). Genes in clusters with high similarity are labeled. (f,g) Eigengene expression of gene modules identified from Weighted correlation network analysis (WGCNA) in cells containing different perturbations relative to unperturbed cells (f). Enriched pathways in each module are shown in g.

We identified differentially expressed genes (DEGs) caused by each perturbation, and determined the similarity of DEGs among all perturbations (see Method for details). This analysis revealed clusters of genes that shared common DEG signatures (Fig. 6e). As expected, knockdown of functionally related genes had similar transcriptomic consequences. For example, knockdown several mitochondria-related genes, namely *COX10, NDUFS8, NDUFV1, MRPL10,* and *SOD2,* resulted in similar DEGs, as did knockdown of the anti-apoptotic genes *BNIP1* and *XIAP.* However, we also identified unexpected gene clusters. For example, knockdown of *VPS54, PAXIP1*, and *PON2* caused highly correlated transcriptomic changes (Fig. 6e, Extended Data Fig. 4c). *VPS54* and PON2 have been implicated in Amyotrophic Lateral Sclerosis (ALS)^82,83^ whereas PAXIP1 is associated with Alzheimer’s disease (AD)^84^. Shared transcriptomic changes included: (i) increased expression of EBF3, an apoptosis inducer^85^ that is also upregulated in the hippocampus of AD model mice^86^, (ii) decreased expression of components of neurofilaments, namely *NEFL, NEFM,* and *NEFH.* Importantly, significantly decreased expression of *NEFL* and *NEFM* was also found in neurons of early-stage AD patients in a recent single-cell transcriptomic study^87^ and (iii) decreased expression of other genes important for neuronal function, including *PCDH11X* and *PCDH11Y,* encoding protocadherin proteins, and *SYT2*, encoding synaptotagmin 2.

To identify genes that were co-regulated under different genetic perturbations, we performed weighted gene co-expression network analysis (WGCNA^88^). This analysis identified 10 modules for CRISPRi and 8 modules for CRISPRa that were co-regulated across different perturbations (Fig. 6f,g). These modules contained genes enriched in various pathways (Fig. 6f). Interestingly, we found a cluster of genes including *INSR, ATP5F1C, SOX5, GSX2* and *GBX2* (Fig. 6e, CRISPRa) overexpression of which downregulated a gene module related to neurogenesis (M1) and upregulated a gene module related to the cell cycle (M5) (Fig. 6g, bottom), suggesting that overexpression of these genes interfered with neuronal differentiation thus kept cells in a proliferating state. This result was expected for *SOX5*, *GSX2* and *GBX2*, which are transcription factors maintaining neural progenitor cell fate and/or their self-renewal^89–91^, but unexpected for *INSR* (encoding the insulin receptor) and *ATP5F1C* (encoding a subunit of mitochondrial ATPsynthase), suggesting these genes might have uncharacterized functions in neuronal fate regulation, which require further investigation.

The CROP-seq data also generated testable hypotheses for disease mechanisms. For example, *NQO1,* encoding the NAD(P)H:Quinone Oxidoreductase 1, is thought to be a cytoprotective factor through its antioxidant functions^92^. Elevated levels and activity of NQO1 have been found in the brains of patients with different neurodegenerative diseases^93–98^, as well as in patient iPSC-derived neurons^98^. The upregulation of NQO1 was proposed to be a neuroprotective mechanism against oxidative stress in neurodegenerative diseases. Paradoxically, however, NQO1 overexpression showed a strong negative impact on neuronal survival in our pooled CRISPRa screen (Fig. 1e). This finding suggests the intriguing hypothesis that elevated NQO1 in the context of neurodegenerative diseases could be neurotoxic, rather than neuroprotective.

Our CROP-Seq data revealed numerous transcriptomic changes induced by NQO1 overexpression (Fig. 6d, Fig. 7a-c), providing several potential hypotheses for the mechanisms underlying NQO1 toxicity.

**Fig. 7:**
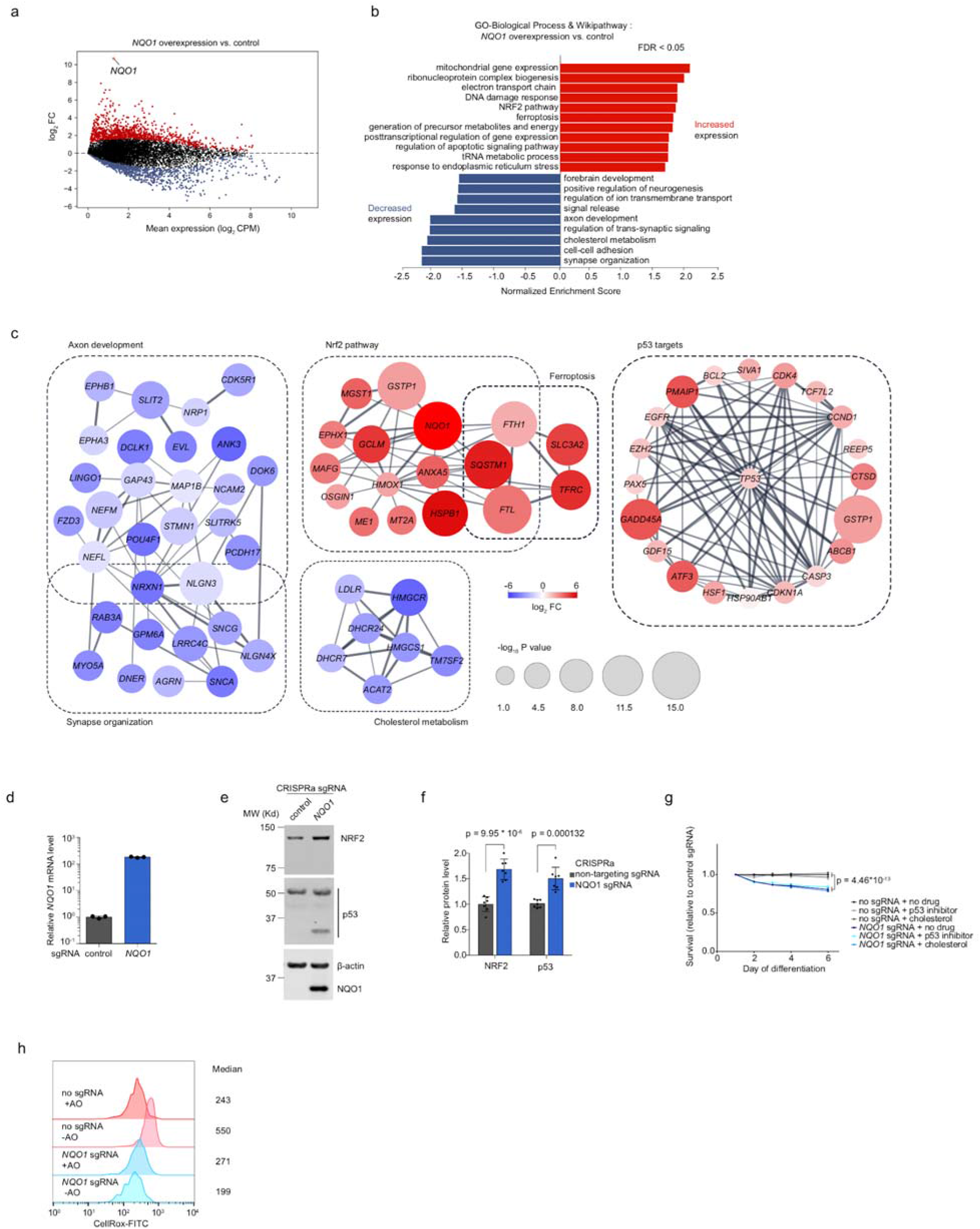
Overexpression of NQO1 induces unexpected transcriptome changes in human iPSC-derived neurons that provide hypotheses for its toxicity. (a) Transcriptomic changes induced by *NQO1* overexpression in neurons. Significantly upregulated or downregulated genes (FDR < 0.01) are shown in red or blue, respectively. (b) Pathway analysis showing enriched pathways in upregulated and downregulated genes in *NQOl*-overexpressing neurons. (c) String-db association networks of selected pathways enriched in upregulated and downregulated genes. Genes with stronger associations are connected by thicker lines. Colors and sizes of nodes reflect log_2_-fold changes (log_2_FCs) and significance (-log_10_P) of differentially expressed genes, respectively. (d) NQO1 overexpression by CRISPRa. qPCR quantification of the relative fold change of NQO1 mRNA levels in CRISPRa-neurons expressing a NQO1 sgRNA as compared to a nontargeting control sgRNA in the presence TMP (mean +/− sd, n = 3 technical replicates). NQO1 levels were normalized to the housekeeping gene ACTB. (e) Western blot showing protein levels of NRF2, p53 and NQO1 in CRISPRa-neurons expressing a NQO1 or non-targeting control sgRNA. ß-actin was used as a loading control. (f) Quantification of NRF2 and p53 levels for CRISPRa-neurons expressing a NQO1 or nontargeting control sgRNA (mean +/− sd, n = 7 independent experiments). P value was calculated using Student’s t-test. (g) Survival of CRISPRa neurons expressing the NQO1 sgRNA relative to non-targeting control sgRNA, quantified by longitudinal imaging for number of BFP+ (sgRNA+) cells on Days 1,2,3,4 and 6 of differentiation. Cells were cultured in the presence of vehicle, 10 μM p53 inhibitor Pifithrin-α hydrobromide or 10 μg/ml cholesterol. Mean +/− sd of 10 replicate wells for each condition at each data point is shown. 4 imaging fields at 20X were taken for each well. P value was calculated using Student’s t-test. (h) ROS levels in CRISPRa-neurons expressing a NQO1 or non-targeting control sgRNA in +AO and -AO conditions, measured by CellRox via flow cytometry. Median signal intensities are indicated.

First, we observed increased expression of many p53 target genes (including p53 itself). A previous study reported that overexpression of NQO1 stabilizes p5 3^99^. The induced p53 activity may contribute to the strong toxic survival phenotype of NQO1 overexpression in neurons. Indeed, we found that p53 protein levels were increased upon NQO1 overexpression (Fig. 7e,f). However, pharmacological inhibition of p53 did not rescue neurons from the toxicity of NQO1 overexpression (Fig. 7g), arguing against p53 as the effector of NQO1 toxicity.

Second, the toxicity of NQO1 overexpression may be caused by other mechanisms, such as downregulation of genes involved in axon development or cholesterol metabolism, which we observed in the CROP-seq data (Fig. 7b,c). Supplementation of culture media with cholesterol did not rescue the neurotoxicity of NQO1 overexpression (Fig. 7g), suggesting that cholesterol deficiency is not the main mechanism of toxicity.

Third, NQO1 overexpression strongly induced target genes of the transcription factor Nrf2 (Fig. 7b,c), and we confirmed that Nrf2 was upregulated at the protein level (Fig. 7e,f). This finding was surprising, because Nrf2 canonically functions upstream of *NQO1* by inducing the expression of many antioxidant genes, including *NQO1,* in response to oxidative stress^100^. We ruled out the possibility that NQO1 overexpression triggered ROS to activate Nrf2 (Fig. 6h), suggesting that NQO1 induces Nrf2 through an unknown mechanism. Intriguingly, the Nrf2 pathway is also induced in some neurodegenerative diseases, both in patient brains^101^ and in patient iPSC-derived neurons^98^. While the concordant upregulation of Nrf2 and NQO1 would canonically be interpreted as a protective response to oxidative stress, with NQO1 as a downstream target of Nrf2, our results suggest that NQO1 overexpression could be upstream of Nrf2, and that the elevation of NQO1 levels in patients could actually contribute to disease, rather than being a protective mechanism.

Taken together, our CROP-seq results provide a rich resource for investigating the consequences of perturbations of neurodegenerative disease-associated genes in human neurons and for generating testable hypotheses for the neurodegenerative disease mechanisms and potential therapeutic strategies.

### CRISPRbrain, an open-access data commons for functional genomics screens in differentiated human cell types

To make the large dataset generated in this study publicly accessible, and to facilitate visualization, interactive exploration, and direct comparison with functional genomics screening data in a broad range of differentiated human cell types generated by many research groups, we developed a data commons, named CRISPRbrain (https://crisprbrain.org) (Fig. 8a).

**Fig. 8:**
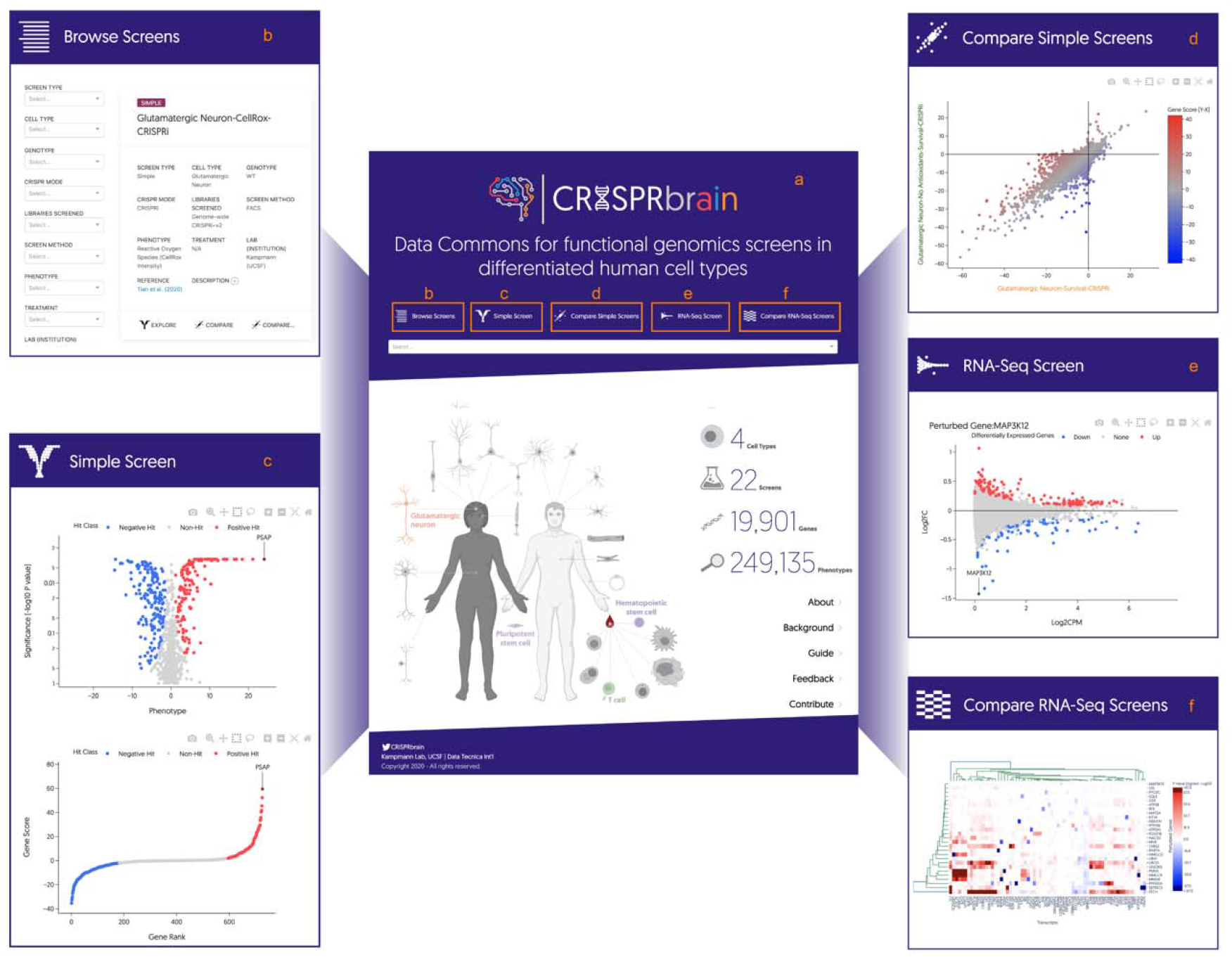
CRISPRbrain, a Data Commons for functional genomics screens in differentiated human cell types. (a) CRISPRbrain (https://crisprbrain.org) is a Data Commons organizing results from genetic screens for different phenotypes in different human cell types, by different research groups. (b) Screens can be browsed and searched based on a range of parameters, and full text search. (c,d) Survival- and FACS-based screens (“simple screens”) can be visualized and explored interactively, and compared pairwise. (e,f) RNA-Seq-based screens can be explored one perturbed gene at a time as MA plots (e), or globally in a hierarchically clustered heatmap (f).

CRISPRbrain currently features screens with “simple” readouts (such as survival and fluorescent reporter levels), as well as screens with complex readouts (such as transcriptomes) Screens can be browsed and searched based on parameters of interest, such as cell type, genotype, mode of CRISPR perturbation, screen method and phenotype, as well as full text search (Fig. 8b).

For simple screens, the phenotypes for the perturbed genes can be displayed as volcano plots and rank plots (Fig. 8c). Data points can be selected individually or in groups to display information about the perturbed gene(s). Gene names can be entered to label them in the graph. Data for the entire screen or for selected genes can be exported for offline analysis. Two screens of interest can be compared in a scatter plot (Fig. 8d).

For RNA-Seq-based screens, genes of interest can be selected to display the transcriptomic phenotype resulting from perturbation of the gene (Fig. 8e). Individual data points can be selected to display information about the transcript, and underlying data can be exported for offline analysis. Alternatively, the results from the entire screen can be explored in a hierarchically clustered heatmap of RNA-Seq phenotypes (Fig. 8f).

We invite all research groups to contribute their functional genomics datasets to CRISPRbrain, with the ultimate goal to build a comprehensive atlas of gene function in all human cell types.

## Discussion

In this study, we developed a CRISPRa platform in human iPSC-derived neurons to enable robust gene overexpression screens, complementing our previously established CRISPRi platform^3^. We demonstrated the power of these complementary screening approaches in multiple large-scale screens in human iPSC-derived neurons.

We determine the first comprehensive inventories of genes that, when depleted or activated, modulate the survival of human neurons. Intriguingly, many of these genes only affect survival in neurons, but not stem cells or cancer cells, supporting the notion that neurons have cell-type specific vulnerabilities, which may explain why defects in some generally expressed genes specifically cause neurological diseases.

Neurons, as one of the longest-living cell types in the human body, are challenged by various stresses in aging and disease. Due to their post-mitotic nature, neurons do not have the ability to ‘self-renew’ by cell division. Therefore, robust stress response mechanisms are required for neurons to maintain long-term health. One of the predominant types of stress in aging and neurodegenerative diseases is oxidative stress^6,7^, which is induced by excessive accumulation of ROS in the cell. The brain is highly susceptible to ROS and ferroptosis, due to its high levels of oxygen consumption, abundant redox-active metals such as iron and copper, limited antioxidants and high levels of PUFAs^102^. A large body of evidence has implicated oxidative stress, iron accumulation and ferroptosis in neurodegenerative diseases^103–105^, yet a comprehensive understanding of how neurons regulate redox homeostasis and maintain survival under oxidative stress is lacking. To this end, we decided to investigate neuronal redox homeostasis and survival under oxidative stress conditions using our CRISPRi/a screening platform.

Compared to other recent genetic screens focusing on oxidative stress toxicity in human cells^106,107^, our screens are unique in the following aspects. First, we used post-mitotic neurons instead of cancer cell lines. Second, we induced milder oxidative stress in cells by prolonged culture of cells in an antioxidant-free medium, compared to severe oxidative stress induced by paraquat or H2O2 in other studies. Third, we screened not only based on cell survival, but also on direct ROS and lipid peroxidation levels, while other studies focused on survival only. Given these major differences, it is not surprising that our screens identified many unique hits not identified in previous screens.

Numerous novel biological insights have emerged from our rich datasets. *GPX4* and genes related to *GPX4* synthesis are indispensable for neurons to survive oxidative stress. Given the major role of *GPX4* in reducing lipid peroxidation and suppressing ferroptosis, this result suggests that lipid peroxidation-induced ferroptosis, rather than other forms of cell death, may be the main cause of neuronal loss under oxidative stress conditions that are commonly found in the brains of patients with neurodegenerative diseases. This is supported by numerous studies reporting high levels of iron and lipid peroxidation in neurodegenerative disease patient brains^108,109^, and by the finding that ferroptosis inhibitors such as iron chelators and the lipid-peroxidation inhibitor Ferrostatin-1 are neuroprotective in animal and cellular models of neurodegenerative diseases^110–113^.

We also found that that disruption of lipid metabolism, in particular glycosphingolipid degradation in the lysosome, by loss of prosaposin (*PSAP*), drives the formation of lipofuscin in neurons, which leads to iron accumulation and strongly induces ROS production, oxidizing lipids and leading to neuronal ferroptosis under oxidative stress. While lipofuscin has traditionally been considered a byproduct of aging and as a consequence of defective cellular homeostasis, its physiological and pathological functions have not been well characterized. This is largely due to the lack of robust systems to model lipofuscin given the fact that lipofuscin is normally only formed in aged post-mitotic cells. Our *PSAP* KO neurons provide a reliable genetic system to model and study the biology of lipofuscin in young, live human neurons. Our result suggests a direct pathogenic role of lipofuscin in inducing neuronal ferroptosis. The accumulation of lipofuscin is a pathological hallmark of many degenerative diseases, such as neuronal ceroid lipofuscinosis (NCL)^114^ and inherited age-related macular degenerations^115,116^. It has also been characterized in Alzheimer’s disease^117,118^, Parkinson’s disease^119^, Hungtinton’s disease^120,121^ and GRN-associated frontotemporal dementia (FTD)^122^. Our findings suggest that inhibiting lipofuscin formation or subsequent ferroptosis may serve as new therapeutic strategies for these diseases.

Our results highlight the importance of lipid homeostasis, in particular balanced levels of glycosphingolipids, for neuronal health. Among all tissues in the human body, the brain is one of the richest in lipid content and lipid diversity. Lipids are not only an essential structural component of membranes, but also important signaling molecules in the brain^123^. Therefore, maintaining lipid homeostasis is of vital importance for brain cells, especially neurons with long neurites and dynamic synaptic vesicles release and recycling. Abnormal lipid metabolism has been observed in neurodegenerative diseases^124–126^. Accumulation of some glycosphingolipids, especially simple gangliosides, caused by inhibition of lysosome membrane recycling, contributes to neurodegeneration both in cultured neurons and in animal models^127^.

Many important questions around neuronal lipid and redox homeostasis and ferroptosis remain to be investigated. For example, since neurons have very long neurites, how do neurons sense and respond to a local lipid peroxidation event on the cell membrane? Are GPX4 or other enzymes recruited to sites of lipid peroxidation, or are peroxidized lipids internalized and delivered to GPX4 or other enzymes for detoxification? What are the sensors and mediators in these processes? How do peroxidized lipids cause death? Is it a passive physical process or a regulated biological program, and are there neuron-specific aspects of ferroptosis? If the latter is true, what are the players mediating cell death downstream of lipid peroxidation? Many of these questions can be readily investigated using our functional genomics platform.

There are several areas for future technology development. First, a robust inducible CRISPRi system that allows temporal control of gene knockdown in mature neurons will help avoiding false-positive screening phenotypes due to interference with the differentiation process. For example, in the CRISPRi survival screen, we unexpectedly identified genes in the N6- methyltransferase writer complex, including *ZC3H13, METTL3,* and *METTL4,* as strong protective hits when knocked down (Fig. 1e). The N6-methyltransferase writer complex is responsible for m6A RNA modification, which regulates a variety of biological processes including neuronal differentiation^128,129^. The inducible CRISPRi system will help to determine whether the strong protective phenotypes of these genes are false positive due to defects in neuronal differentiation. Second, we have previously used our platform for arrayed high-content imaging screens for complex neuronal phenotypes, such as neurite outgrowth^3^. Such imaging-based screens could be applied to study other neuronal phenotypes, such as electrophysiological signals (by voltage imaging or calcium imaging), synaptic vesicle dynamics and axonal transport, and cell non-autonomous phenotypes such as glia-neuron interaction. Imaging phenotyping can also be coupled with simultaneous transcriptomic profiling, thus allowing the association of gene expression with imaging phenotypes for a given genetic perturbation. Lastly, by combining our platform with base editor or prime editing technology^130^, we will be able to directly assess the effect of disease-associated mutations in human neurons on various cellular phenotypes in a scalable, massively parallel format. While base editor screens to assess human variants have been recently conducted in cancer cell lines^131^, it is of great interest to investigate how variants associated with different human diseases could affect various cell types differently given the fact that a lot of human diseases are tissue-specific.

In the past decade, tremendous efforts have advanced the molecular characterization of various human cell types. For example, through global collaboration in the Human Cell Atlas project, gene expression profiles for >30 human organs in > 4.5M cells have been obtained so far (https://data.humancellatlas.org/). Beyond gene expression, characterizing gene function in different cell types will be the next step towards building complete reference maps for the human body at the cellular level. We anticipate that our iPSC-based functional genomics platforms can be broadly applied to a variety of human differentiated cell types. Our CRISPRbrain Data Commons is complementary to the Human Cell Atlas and serves as an open-access platform for sharing and exploring functional genomics screening data in differentiated human cell types across labs. In contrast to the Cancer Dependency Map^132^, BioGRID ORCS^133^ and CRISP-view^134^, CRISPRbrain is the first database focused on screens in non-cancerous human cell types for a broad range of phenotypes, including high-dimensional transcriptomic phenotypes. Future versions of CRISPRbrain will support other high-dimensional phenotypes, such as imaging and electrophysiological data, and enable cloud computing and machine learning, thus providing researchers with a suite of tools to process, visualize, analyze and contribute functional genomics screening data in different human cell types. Parallel genetic screens across the full gamut of isogenic human cell types will uncover context-specific roles of human genes, leading to a deeper mechanistic understanding of how they control human biology and disease.

## Methods

### HEK293 culture

HEK293 cells were cultured in DMEM (GIBCO; Cat. No. 10313-039) supplemented with 10% FBS and 1% penicillin/streptomycin. For CRISPRi knockdown, sgRNAs were introduced into HEK293 cells with CRISPRi machinery^135^ via lentiviral delivery. Cells were selected by 2ug/ml puromycin for 2 days and recovered for 2 days. Phenotypes were evaluated 5 days after infection.

### Human iPSCs culture and neuronal differentiation

Human iPSCs from WTC11 and NCRM5 backgrounds^3^ were cultured in StemFlex Medium (GIBCO/Thermo Fisher Scientific; Cat. No. A3349401) in plates or dishes coated with Growth Factor Reduced, Phenol Red-Free, LDEV-Free Matrigel Basement Membrane Matrix (Corning; Cat. No. 356231) diluted 1:100 in Knockout DMEM (GIBCO/ v; Cat. No. 10829-018). StemFlex Medium was replaced every other day or every day once cells reached 50% confluence. When 80%-90% confluent, cells were dissociated using StemPro Accutase Cell Dissociation Reagent (GIBCO/Thermo Fisher Scientific; Cat. No. A11105-01) at 37°C for 5 min, centrifuged at 200 g for 5 min, resuspended in StemFlex Medium supplemented with 10 nM Y-27632 dihydrochloride ROCK inhibitor (Tocris; Cat. No. 125410) and plated onto Matrigel-coated plates or dishes at desired number. Studies with human iPSCs at UCSF were approved by the Human Gamete, Embryo, and Stem Cell Research (GESCR) Committee.

For individual gene knockdown or overexpression in CRISPRi- or CRISPRa-iPSCs, sgRNAs were introduced into iPSCs via lentiviral delivery. Cells were selected by 2 μg/ml puromycin for 2-4 days and recovered for 2 days. Phenotypes were evaluated 5-7 days after infection.

The CRISPRi- and CRISPRa-iPSC lines used in this study were engineered to express mNGN2 under a doxycycline-inducible system in the AAVS1 safe harbor locus. For their neuronal differentiation, we followed our previously described protocol^3^. Briefly, iPSCs were predifferentiated in matrigel-coated plates or dishes in N2 Pre-Differentiation Medium containing the following: Knockout DMEM/F12 (GIBCO/Thermo Fisher Scientific; Cat. No. 12660-012) as the base, 1X MEM Non-Essential Amino Acids (GIBCO/Thermo Fisher Scientific; Cat. No. 11140-050), 1X N2 Supplement (GIBCO/ Thermo Fisher Scientific; Cat. No. 17502-048), 10 ng/mL NT-3 (PeproTech; Cat. No. 450-03), 10ng/mL BDNF (PeproTech; Cat. No. 450-02), 1 μg/mL Mouse Laminin (Thermo Fisher Scientific; Cat. No. 23017-015), 10nM ROCK inhibitor, and 2 μg/mL doxycycline to induce expression of mNGN2. After three days, on the day referred to hereafter as Day 0, pre-differentiated cells were re-plated into BioCoat Poly-D-Lysine-coated plates or dishes (Corning; assorted Cat. No.) in regular Neuronal Medium, which we will refer to as +AO Neuronal Medium, containing the following: half DMEM/F12 (GIBCO/Thermo Fisher Scientific; Cat. No. 11320-033) and half Neurobasal-A (GIBCO/Thermo Fisher Scientific; Cat. No. 10888-022) as the base, 1X MEM Non-Essential Amino Acids, 0.5X GlutaMAX Supplement (GIBCO/Thermo Fisher Scientific; Cat. No. 35050-061), 0.5X N2 Supplement, 0.5X B27 Supplement (GIBCO/Thermo Fisher Scientific; Cat. No. 17504-044), 10ng/mL NT-3, 10ng/mL BDNF and 1 μg/mL Mouse Laminin. For -AO experiments, we used a medium we refer to as -AO Neuronal Medium, in which B-27 Supplement minus antioxidants (GIBCO/Thermo Fisher Scientific; Cat. No. 10889-038) was used instead of regular B27 in the +AO Neuronal Medium. Neuronal Medium was half-replaced every week.

### Generation of iPSC-derived neural progenitor cells and astrocytes

WTc11 iPSCs with CRISPRi machinery stably integrated into CLYBL locus were differentiated into NPCs using an embryoid body-based dual SMAD inhibition neural induction protocol as described in Topol et al.^137^ The NPCs were then sorted for CD133+/CD271- cells as described in Cheng et al.^138^ to remove contaminating neural crest cells. The sorted NPCs were then expanded and differentiated to astrocytes using the protocol described in TCW et al.^139^.

For CRISPRi knockdown, sgRNAs were introduced into NPCs and astrocytes via lentiviral delivery. Cells were selected by 2ug/ml puromycin for 2-5 days and recovered for >2 days. Phenotypes were evaluated 5-10 days after infection.

### Microglia differentiation

iPSC-Microglia were differentiated from CRISPRi iPSCs using published protocols^140^ with minor modifications. Briefly, iPSCs cultured in StemFlex media with colonies at 60-80% confluency were dissociated to single cells with Accutase, collected and plated at 10,000 per well in 96-well ultra-low attachment plates in 100 mL embryoid body medium (10 mM ROCK inhibitor, 50 ng/mL BMP-4 (Peprotech; Cat. No. 120-05), 20 ng/mL SCF (Peprotech; Cat. No. 300-07), and 50 ng/mL VEGF (Peprotech; Cat. No. 100-20) in E8 (Gibco; A1517001)), before centrifugation at 300g for 3 min. Embryoid bodies were cultured for 4 days, with a half medium change after 2 days. At day 4, embryoid bodies were carefully collected and transferred into a 15 ml Eppendorf tube, and left to settle. The embryoid medium was aspirated and 15 to 20 embryoid bodies were plated per well in 6-well plates and cultured in 3 mL hematopoetic medium (2 mM GlutaMax, 1x Antibiotic-Antimycotic (Thermo Fisher Scientific; Cat. No. 15240062), 55 mM 2-mercaptoethanol (BioRad; Cat. No. 1610710), 100 ng/mL M-CSF (Peprotech; Cat. No. 300-25), and 25 ng/mL IL-3 (Peprotech; Cat. No. 200-03) in X-VIVO 15 (Lonza; Cat. No. BE02-060F). Two thirds of media were exchanged every 3-4 days. Microglia progenitors were harvested from suspension after 14-21 days and plated onto PDL-coated plates in microglia maturation media (2 mM GlutaMax, 1x Antibiotic-Antimycotic, 100 ng/mL IL-34 (Peprotech; Cat. No. 200-34), and 10 ng/mL GM-CSF (Peprotech; Cat. No. 300-03) in Advanced RPMI (Gibco, Cat. No. 12633012)). Microglia progenitors were further differentiated for 8 days with full medium change every 2-3 days before using them for experiments.

For CRISPRi knockdown, sgRNAs were introduced into iPSCs via lentiviral delivery. After 2-4 days of puromycin selection and 2 days of recovery, these iPSCs were differentiated into microglia following the protocol as described above. Phenotypes were evaluated after 10 days of microglia differentiation.

### Molecular cloning

The CLYBL-targeting inducible CRISPRa vector pRT43 containing CAG-driven DHFR-dCas9- VPH-T2A-EGFP was generated by sub-cloning DHFR-dCas9-VPH-T2A-EGFP from plasmid PB-CAG-DDdCas9VPH-T2A-GFP-IRES-Neo to the downstream of a CAG promoter in CLYBL-targeting plasmid pUCM-CLYBL-hNIL digested by SalI and EcoRV. (PB-CAG- DDdCas9VPH-T2A-GFP-IRES-Neo was a gift from Timo Otonkoski (Addgene plasmid # 102886; http://n2t.net/addgene:102886; RRID:Addgene_102886) and pUCM-CLYBL-hNIL was a gift from Michael Ward (Addgene plasmid # 105841; http://n2t.net/addgene:105841; RRID:Addgene_105841))

### CRISPRa-iPSC cell line generation

WTC11 iPSCs harboring a single-copy of doxycycline-inducible mouse NGN2 at the AAVS1 locus (Ngn2-iPSCs,^9^) were used as the parental iPSC line for further genetic engineering. iPSCs were transfected with pRT43 containing DHFR-dCas9-VPH and TALENS targeting the human CLYBL intragenic safe harbor locus (between exons 2 and 3) (pZT-C13-R1 and pZT- C13-L1, gifts from Jizhong Zou (Addgene plasmid # 62196; http://n2t.net/addgene:62196; RRID:Addgene_62196, and Addgene plasmid # 62197; http://n2t.net/addgene:62197; RRID:Addgene_62197) using Lipofectamine Stem (Invitrogen/Thermo Fisher Scientific; Cat. No. STEM00003). Monoclonal lines were isolated by limiting dilution and CLYBL integration was confirmed by PCR genotyping. Karyotype testing (Cell Line Genetics) was normal for the clonal line used for further experiments in this study, which we termed CRISPRa iPSCs.

### Genome-wide survival-based and FACS-based screens

The genome-wide CRISPRi and CRISPRa libraries hCRISPRi-v2 and hCRISPRa-v2^10^, consisting of 7 sublibraries each (H1-H7), were packaged into lentivirus as previously described^3^. CRISPRi- and CRISPRa-iPSCs were infected by the sgRNA libraries at MOIs of 0.4-0.6 (as measured by the BFP fluorescence from the lentiviral vector) with approximately 1000x coverage per library element. Two days after infection, the cells were selected for lentiviral integration using puromycin (1 μg/mL) for 3 days as the cultures were expanded for the screens. After selection and expansion, a fraction of the cells (Day −3 iPSCs) were harvested and subjected to sample preparation for next-generation sequencing. Another fraction of Day −3 iPSCs, with a cell count corresponding to 1000x coverage per library element, were differentiated into neurons as described in the Human iPSCs Culture and Neuronal Differentiation subsection.

Neurons were cultured in either the +AO or -AO Neuronal Medium (see Human iPSCs Culture and Neuronal Differentiation subsection) for ten days. For the survival screens, Day 10 neurons were harvested and subjected to sample preparation for next-generation sequencing. For the FACS screens, Day 10 CRISPRi neurons cultured in the +AO medium and -AO medium were dissociated using Papain (Worthington; Code: PAP2; Cat. No.LK003178) and stained by CellRox Green (Invitrogen/Thermo Fisher Scientific; Cat. No. C10444) and Liperfluo (Dojindo Molecular Technologies, Inc.; Cat. No. L248-10) respectively (see Cell Staining by Fluorescent Probes) and sorted into high and low signal populations in each screen corresponding to the top 40% and the bottom 40% of the staining signal distribution, followed by sample preparation for next-generation sequencing.

For each screen sample, genomic DNA was isolated using a Macherey-Nagel Blood L kit (Macherey-Nagel; Cat. No. 740954.20). sgRNA-encoding regions were amplified and sequenced on an Illumina HiSeq- 4000 as previously described^4^.

### Focused secondary screens

The focused secondary screen library contained 2,190 sgRNAs targeting 730 genes that were hits in at least one of the ROS and lipid peroxidation screens with 3 sgRNAs per gene selected based on their phenotypes in the primary screens, and 100 non-targeting control sgRNAs (Supplementary Table 4). A pool of sgRNA-containing oligonucleotides were synthesized by Agilent Technologies and cloned into our optimized sgRNA expression vector as previously described^4^. CRISPRi-iPSCs were transduced with the batch characterization library, puromycin selected, and differentiated into neurons as for the primary screens. Day10 neurons were stained with FeRhoNox-1 (Goryo Chemical; Cat. No. GC901) or Lysotracker-green (Cell Signaling Technology; Cat. No. 8783S) and sorted into high and low signal populations corresponding to the top 40% and bottom 40% of the staining signal distribution. Screen samples were processed and sequenced by next-generation sequencing as described above.

### CROP-seq

For the CROP-seq experiments, we included 184 genes for CRISPRi and 100 genes for CRISPRa, which were hits in at least one of our genome-wide pooled screens and were also associated with human neurodegenerative diseases. We curated a list of genes associated with neurodegenerative diseases based on the literature and the DisGeNET database (https://www.disgenet.org/). The CROP-seq libraries included 2 sgRNAs per gene plus 6 nontargeting control sgRNAs, for a total of 374 sgRNAs for CRISPRi and 206 sgRNAs for CRISPRa (Supplementary Table 4). Top and bottom strands of sgRNA oligos were synthesized (Integrated DNA Technologies) and annealed in an arrayed format. The annealed sgRNAs were then pooled in equal amounts and ligated into our optimized CROP-seq vector^3^.

The CROP-seq experiments were carried out similarly as previously described^3^. Briefly, Day 0 CRISPRi and CRISPRa neurons were infected by the corresponding CROP-seq sgRNA library at a MOI of 0.1-0.2, followed by puromycin selection at 4 μg/ml for 3 days and recovery. On Day 10, neurons were dissociated with Papain and approximately 98,000 CRISPRi neurons and 50,000 CRISPRa neurons were loaded into 10X chips with about 25,000 input cells per lane. Sample preparations were performed using the ChromiumNext GEM Single Cell 3 ? Reagent Kits v3.1 (10X Genomics; Cat. No.PN-1000121) according to the manufacturer’s protocol. To facilitate sgRNA assignment, sgRNA-containing transcripts were additionally amplified by hemi-nested PCR reactions as described^3^. The sgRNA-enrichment libraries were separately indexed and sequenced as spike-ins alongside the whole- transcriptome scRNA-seq libraries using a NovaSeq 6000 using the following configuration: Read 1: 28, i7 index: 8, i5 index: 0, Read 2: 91.

### Cell Staining by Fluorescent Probes

All stains were performed according to manufacturing protocols. For Filipin staining, cells were washed with PBS three times and fixed with 3% paraformaldehyde for 1 hr at room temperature. Cells were then washed 3 times with PBS and incubated with 0.05 mg/ml Filipin III from Streptomyces filipinensis (Sigma; Cat. No. F4767-1MG) in PBS for 2 h at room temperature. Cells were washed with PBS 3 times before analysis. For staining using other live cell probes, cells were washed with PBS and incubated in DMEM containing appropriate concentrations of the probes at 37 °C as detailed below. Cells were washed with PBS before analysis. Concentrations and staining conditions for different probes were as follows: CellRox Green, 2.5 μM for 30 minutes; Lysotracker Green, 50 nM for 5 minutes; FeRhoNox-1, 5 μM for 60 minutes; Liperfluo, 5 μM for 30 minutes; C11-BODIPY, 2.5 μM for 30 minutes; FerroOrange (Dojindo Molecular Technologies, Inc.; Cat. No. F374-10), 1 μM for 30 minutes; Hoechst 33342 (Thermo Fisher Scientific; Cat. No. H3570), 1 μg/ml for 10 minutes; Propidium Iodide (PI)(Thermo Fisher Scientific; Cat. No. P1304MP), 1 μg/ml for 10 minutes.

### Immunofluorescence

Cells were washed with PBS and fixed with 4% paraformaldehyde for 15 min at room temperature. After washing with PBS for 3 times, cells were permeabilized with 0.1% Triton X-100 for 10 min and blocked with 5% normal goat serum with 0.01% Triton X-100 in PBS for 1 hr at room temperature. Cells were then incubated with primary antibodies diluted in blocking buffer at 4 °C overnight. After that, cells were washed with PBS for 3 times and incubated with secondary antibodies diluted in blocking buffer for 1 hr at room temperature. Cells were then washed with PBS for 3 times and stained with 10 μg/ml Hoechst 33342 (Thermo Fisher Scientific; Cat. No. H3570) for 10 min. Cells were imaged using a confocal microscope (Leica SP8) or an IN Cell Analyzer 6000 (GE; Cat. No. 28-9938-51). Primary antibodies used for immunofluorescence in this study were as follows: rabbit anti-*PSAP* antibody (1:50 dilution; Proteintech; Cat. No. 10801-1-AP), mouse anti-LAMP2 antibody (1:100 dilution; abcam; Cat. No. ab25631), rabbit anti-GM1 antibody (1:20 dilution; abcam; Cat. No. ab23943), rabbit anti- LC3B antibody (1:200 dilution; Cell Signaling Technology; Cat. No. 2775S), chicken anti-TUJ1 antibody (1:500; AVES; Cat. No. TUJ), mouse anti-S100β antibody (1:500; Sigma; Cat. No. S2532), mouse anti-Nestin antibody (1:500; Sigma; Cat. No. MAB5326) and mouse anti-GPR34 antibody (1:50; R&D Systems; Cat. No. MAB4617). Secondary antibodies used in this study were as follows: goat anti-rabbit IgG Alexa Fluor 555 (1:500 dilution; abcam; Cat. No. ab150078), goat anti-mouse IgG Alexa Fluor 488 (1:500 dilution; abcam; Cat. No. ab150113) and goat anti-chicken IgG Alexa Fluor 647 (1:500 dilution; abcam; Cat. No. ab150171).

### Drug treatments

Drugs (Z-VAD-FMK (Sigma-Aldrich; Cat. No. V116), Ferrostatin-1 (Sigma-Aldrich; Cat. SML0583), DFO (Sigma-Aldrich; Cat. No. D9533), Pifithrin-α hydrobromide (Tocris; Cat. No. 1267) and cholesterol (complexed with MβCD; Sigma-Aldrich; Cat. No. C4951)) were dissolved in DMSO or water as per manufacturer’s instructions. Drug concentrations were determined according to previous reports. Specifically, 1μM and 10 μM of Z-VAD-FMK^141^, 1μM and 10 μM of Ferrostatin-1^141^, 10 μM DFO^141^, 10 μM Pifithrin-α hydrobromide^142^ and 10 μg/ml cholesterol^143^ were used.

### Western blots

Cells were lysed in RIPA buffer and 20-30 μg of total proteins were loaded into NuPAGE 4%- 12% Bis-Tris Gels (Invitrogen, Cat# NP0336BOX). Subsequently, the gels were transferred onto nitrocellulose membranes and the membranes were blocked by Odyssey Blocking Buffer (PBS) (LI-COR, Cat#927-50000), followed by overnight incubation with primary antibodies at 4°C. After incubation, the membranes were washed three times with TBST and then incubated with secondary antibodies at room temperature for 1 hr. The membranes were then washed 3 times with TBST and once with TBS and imaged on the Odyssey Fc Imaging system (LI-COR Cat# 2800). Digital images were processed and analyzed using ImageJ.

Primary antibodies used were mouse anti-β-Actin antibody (1:2000 dilution; Cell Signaling Technology; Cat. No. 3700), rabbit anti-*PSAP* antibody (1:1000 dilution; Proteintech; Cat. No. 10801-1-AP) and rabbit anti-LC3B antibody (1:1000 dilution; Cell Signaling Technology; Cat. No. 2775S). Secondary antibodies were IRDye 680RD goat anti-mouse IgG (1:20,000 dilution; LI-COR; Cat. No. 926-68070) and IRDye 800CW goat anti-rabbit IgG(1:20,000 dilution; LI-COR; Cat. No. 926-32211).

### Generating the PSAP KO iPSC line

An sgRNA targeting *PSAP* exon 2 (sgRNA sequence: GGACTGAAAGAATGCACCA) was cloned into plasmid px330-mcherry (px330-mcherry was a gift from Jinsong Li (Addgene plasmid # 98750; http://n2t.net/addgene:98750; RRID:Addgene_98750)). The plasmid was transfected into WT Ngn2-iPSCs using Lipofectamine Stem (Invitrogen/Thermo Fisher Scientific; Cat. No. STEM00003). Monoclonal lines were isolated in 96-well plates by limiting dilution. One clonal line was selected and frameshift indels were confirmed by Sanger sequencing. Protein level KO was confirmed by western blot and immunofluorescence (Fig. 5c,d). A normal karyotype was confirmed (Cell Line Genetics).

### Bulk RNA sequencing

RNA was extracted from cells using the Quick-RNA Miniprep Kit (Zymo; Cat. No. R1054), and 3’-tag RNA-seq was performed by the DNA Technologies and Expression Analysis Core at the UC Davis Genome Center.

### Electron Microscopy

Neurons grown on a poly-D-lysine coated 35-mm ibidi μ-Dish (ibidi; Cat. No. 81156) were fixed with 2.5% glutaraldehyde in 0.1 M sodium cacodylate buffer (EMS) for at least an hour at room temperature. Samples were further post-fixed with 1% osmium tetroxide and 1.6% potassium ferrocyanide, later dehydrated in graded series of ethanols, and embedded in epon araldite resin. Samples were then trimmed, 70nm sections were cut using Ultra cut E (Leica) and stained with 2% uranyl acetate and Reynold’s lead citrate. Images were acquired on a FEI Tecnai 12 120KV TEM (FEI) and data was recorded using UltraScan 1000 Digital Micrograph 3 software (Gatan Inc.)

### STORM super-resolution microscopy

Sample preparation and STORM imaging were performed as described previously^144^. Samples were fixed in 3% paraformaldehyde and 0.1% glutaraldehyde (Electron Microscopy Sciences) in PBS (Corning) for 30 minutes at room temperature, followed by reduction with 0.1% NaBH_4_ in PBS for 10 minutes and three times of washing with PBS. Prior to immunostaining, samples were treated by a blocking buffer (BB) of 3% bovine serum albumin (Sigma) and 0.1% Triton- X100 (Sigma) in PBS for 1 hour at room temperature. Primary antibodies were diluted in BB and labeled at 4C overnight. Unbound antibodies were rinsed three times, 10 minutes each time with a washing buffer (WB) prepared by 10X dilution of BB in PBS. Secondary antibodies Dye-labeled secondary antibodies from Invitrogen or Jackson ImmunoResearch Laboratories were diluted in BB and labeled at room temperature for 1 hour, followed by three times of washing with WB.

Before STORM imaging, the sample was mounted in a standard STORM imaging buffer of 5% [w/v] glucose (Sigma), 0.1 M cysteamine (Sigma), 0.8 mg/mL glucose oxidase (Sigma), and 40 mg/mL catalase (Sigma) in 0.1 M Tris-HCl pH 7.5 (Corning). Two-color STORM was carried out on a custom setup modified from a Nikon Eclipse Ti-E inverted fluorescence microscope^145^ by sequentially imaging the Alexa Fluor 647-labeled GM1 and the CF568-labeled LAMP2 using 647 nm and 560 nm excitations, respectively. A strong (~2 kW/cm^2^) excitation laser (647 nm or 560 nm) was applied to photoswitch most of the AF647 or CF568 dye molecules into the dark state while also exciting the remaining dye molecules at a low density for single-molecule localization. A weak (0-1 W/cm^2^) 405-nm laser was used simultaneously with the 647-nm or 560-nm laser to reactivate dye molecules in the dark state into the emitting state to acquire adequate sampling of the labeled molecules. Images were collected at 110 frames per second with an electron multiplying charge-coupled device camera (Andor iXon Ultra 897) for ~50,000 frames per image.

### Untargeted Lipidomics

The untargeted lipidomics experiment and primary analysis were performed by Cayman Chemical. Briefly, lipids were extracted using a methyl□tert□butyl ether (MTBE)□based liquid liquid method. Cell pellets (approximately 100 μL in volume) were thawed on ice and transferred into 8-mL screw-cap tubes before adding 600 μL MeOH, the 600 μL MeOH containing 200 ng each of the internal standards TG(15:0/18:1□d7/15:0), PC(15:0/18:1□d7), PE(15:0/18:1□d7), PG(15:0/18:1□d7), and PI(15:0/18:1□d7) (EquiSPLASH, Avanti Polar Lipids), and finally 4 mL MTBE. After vigorous vortexing, the samples were incubated at room temperature on a shaker for 1 h. For phase separation, 1 mL water was added, and samples were vortexed and centrifuged for 10 min at 1000 x g. The upper organic phase of each sample was carefully removed using a Pasteur pipette and transferred into a pre weighed empty glass tube. The remaining aqueous phase was re-extracted with 2 mL of clean MTBE/methanol/water 10:3:2.5 (v/v/v). The two upper organic phases were combined and dried under vacuum in a SpeedVac concentrator. The dried lipid extracts were weighed and resuspended in 100 μL isopropanol/acetonitrile 1:1 (v/v) for untargeted lipidomic analysis by LC□MS/MS. Triplicates of samples for WT and *PSAP* KO neurons were analyzed.

### Data analysis

#### Pooled CRISPR screens

Pooled CRISPR screens were analyzed using the MAGeCK-iNC pipeline as described^3^. Briefly, raw sequencing reads from next-generation sequencing were cropped and aligned to the reference using Bowtie^146^ to determine sgRNA counts in each sample. Counts files for samples subject to comparison were entered into the MAGeCK-iNC software^3^ (kampmannlab.ucsf.edu/mageck-inc). Raw Phenotype scores and significance P values were calculated using MAGeCK-iNC for target genes, as well as for ‘negative-control-quasi-genes’ that were generated by random sampling of 5 with replacement from non-targeting control sgRNAs.

The final Phenotype score for each gene was calculated by subtracting the raw Phenotype score by the median of ‘negative-control-quasi-genes’ and then dividing it by the standard deviation of ‘negative-control-quasi-genes’.

A ‘Gene score’ was defined as the product of Phenotype score and -log10(Pvalue). Hit genes were determined based on Gene score cutoff corresponding to an empirical false discovery rate (FDR) of 5%.

#### CROP-seq

CROP-seq analysis was performed similarly to previously described^3^. Cellranger (version 3.1.0,10X Genomics) with default parameters was used to align reads and generate digital expression matrices from single-cell sequencing data.

Approximately 58,000 CRISPRi neurons and 38,000 CRISPRa neurons were detected. The mean reads per cell was around 48,000 for CRISPRi and 36,000 for CRISPRa. The median number of genes detected per cell was around 4,341 for CRISPRi and 3,100 for CRISPRa.

sgRNA-enrichment libraries were analyzed using methods previously described^147^ to obtain sgRNA UMI counts for each cell barcode. For a given cell, sgRNA(s) whose UMI counts were greater than 4 standard deviations of the mean UMI counts of all sgRNAs were assigned to that cell as its identity. Single sgRNAs could be assigned to about 35,000 CRISPRi cells and about 21,000 CRISPRa cells, which were retained for further analysis (Supplementary Table 5)

The Scanpy package^148^ (version 1.4.6) implemented in Python was used for downstream analysis of the digital expression matrices with mapped sgRNA identities. To ensure data quality, a stringent criterion was applied to filter cells based on the number of genes detected (> 2000 for CRISPRi and > 1500 for CRISPRa) and percentage of mitochondrial transcript counts (< 0.15%). Genes that had less than 0.5 UMIs on average in all perturbation groups were filtered out.

Within the population of cells expressing sgRNAs for a specific target gene, the levels of target knockdown or overexpression were heterogeneous in some cases. This could be due to different efficiencies of the two sgRNAs targeting that gene (see Extended Data Fig. 4a for an example), misassignment of sgRNA identities for some cells or stochastic silencing of the CRISPR machinery in some cells. To select cells in which functional perturbations happened, we leveraged an unsupervised outlier detection method based on the local outlier factor (LOF) using the LocalOutlierFactor function in the Python package scikit-learn (version 0.23.0). A similar strategy was described previously^79^. Specifically, for every target gene, we selected two populations of cells, including one population of cells that were mapped with sgRNAs targeting that gene (i.e. perturbed group) and another population of cells that were mapped with nontargeting control sgRNAs (i.e. control group). We then identified differentially expressed genes (DEGs) between the two populations by t-tests at p < 0.05. A gene-expression matrix containing only expression of DEGs in cells from the two populations was generated and principal component analysis (PCA) was performed to reduce the matrix to 4 dimensions. Cells in the control group in the 4-dimensional space were used as the training set to fit a LocalOutlierFactor model. Then, the model was used to determine whether a cell in the perturbed group was an ‘outlier’ based on the extent it deviated from the controls. The ‘outliers’ were considered as cells in which functional perturbations occurred and were retained for downstream analysis. This classification method was particularly useful for CRISPRi when the basal expression level of the target gene was too low to be detected by single-cell RNA-seq thus did not allow selecting cells based on knockdown level of target gene (see Extended Data Fig. 4b for an example).

DEGs for each perturbation group compared to the control group were then determined by t-tests using the diffxpy package in Python.

A mean gene-expression matrix was generated for different perturbation groups (including the control group) by averaging the normalized expression profile of all cells within that group. This matrix was used for weighted correlation network analysis (WGCNA) using the WGCNA package^88^ (version 1.69) implemented in R. The blockwiseModules function was used to detect gene modules that were co-regulated in different perturbation groups and to determine eigengene expression of each module in each perturbation group. Relative eigengene expression values were calculated by subtracting the eigengene expression value of each module in the control group from that in each perturbation group.

#### RNA-seq

Raw sequencing reads from 3 -tag RNA-seq were mapped to the human reference transcriptome (GRCh38, Ensembl Release 97) using Salmon^149^ (v.0.14.139) with the ‘-noLengthCorrection’ option to obtain transcript abundance counts. Gene-level count estimates were obtained using tximport^150^ (v.1.8.040) with default settings. Subsequently, genes with more than 10 counts were retained for differential gene-expression analysis, and adjusted P values (P_adj_) were calculated using DESeq2^151^ (v.1.20.041).

#### Pathway enrichment analysis

Gene Ontology (GO) term and Wikipathways enrichment analysis was performed using WebGestalt (WEB-based Gene SeT AnaLysis Toolkit) using the over-representation analysis (ORA) method^152^.

#### Lipidomics

Lipostar software (Molecular Discovery) was used for feature detection, noise and artifact reduction, alignment, normalization, and lipid identification. For each lipid, the log2-fold change and a significance P value by t-test were determined by comparing the abundances of that lipid in WT and *PSAP* KO neurons (samples in triplicates). The Benjamini-Hochberg (BH) method was used to correct for multiple hypothesis testing.

### Data commons development

CRISPRbrain (https://crisprbrain.org) was developed as an open-access and cloud-based platform striving to make data and code easily accessible to the scientific community. To enable scientific transparency and replication, each dataset in CRISPRbrain is licensed under a Creative Commons Attribution 4.0 International License (CC BY 4.0). To handle a large volume of storage and computation demand, CRISPRbrain is deployed on the cloud with elastic resource allocation. We have prioritized a low-latency interactive user experience by pre-computing and caching complex queries. CRISPRbrain is implemented in a manner that is “future-proof”, supporting scalability and addition of more complex features.

## Author contributions

R.T. and M.K. conceived this study and wrote the manuscript with input from the other authors. R.T. designed and conducted experiments with help from A.A. and J.H. and guidance from M.K.. R.T. performed data analyses. R.Y. performed STORM imaging with guidance from K.X.. N.D. generated iPSC-derived microglia and K.L. generated iPSC-derived astrocytes. S.H.H., M.A.N. and F.F. developed the CRISPRbrain Data Commons with critical input from R.T. and M.K. and feedback from A.B.S.. All authors reviewed and approved the final manuscript.

## Acknowledgements

We thank Li Gan, Bruce Conklin for support and advice; Paul Kennedy and Anna Nummy at Cayman Chemical for untargeted lipidomics; Eric Chow (UCSF), Derek Bogdanoff (UCSF), Angela Detweiler (CZI Biohub), Norma Neff (CZI Biohub) and Michelle Tan (CZI Biohub) for next-generation sequencing; Avi Samelson and Xiaoyan Guo for comments on this manuscript; and Kun Leng, Emmy Li and James Olzmann for discussions. We thank the staff at the University of California Berkeley Electron Microscope Laboratory for advice and assistance in electron microscopy sample preparation and data collection. This research was supported by the Intramural Research Program of the NIH/NINDS, an NIH Director’s New Innovator Award (NIH/ NIGMS DP2 GM119139 to M.K.), NIH/NIA grants (R01 AG062359 and R56 AG057528 to M.K., F30 AG066418 to K.L.), the NINDS Tau Center Without Walls (NIH/NINDS U54 NS100717 to M.K.), an Allen Distinguished Investigator Award to M.K. (Paul G. Allen Family Foundation, to M.K.), a Chan-Zuckerberg Biohub Investigator Award (to M.K.), and a Tau Consortium Investigator Award (Rainwater Charitable Foundation, to M.K.). K.X. is a Chan Zuckerberg Biohub investigator and acknowledges support from the National Institute of General Medical Sciences of the National Institutes of Health (DP2GM132681). A. B. S.’s participation was supported in part by the Intramural Research Program of the National Institute on Aging, National Institutes of Health, part of the Department of Health and Human Services (project number ZIA AG000957-16). CRISPRbrain development was supported in part by a collaboration between the Kampmann Lab, UCSF and Data Tecnica International, LLC.

## Declaration of interests

M.K. has filed a patent application related to CRISPRi and CRISPRa screening (PCT/US15/40449) and serves on the Scientific Advisory Board of Engine Biosciences, Casma Therapeutics, and Cajal Neuroscience.

## Data availability

All screen datasets are publicly available on the CRISPRbrain website (http://crisprbrain.org/). The accession number for the RNA sequencing datasets reported in this paper is GEO: GSE152988 (https://www.ncbi.nlm.nih.gov/geo/query/acc.cgi?acc=GSE152988).

## Code availability

All data analyses were performed using published computational pipelines and standard python/R packages as described in the Methods section. Original codes are available upon request.

---

**Extended Data Fig. 1:**
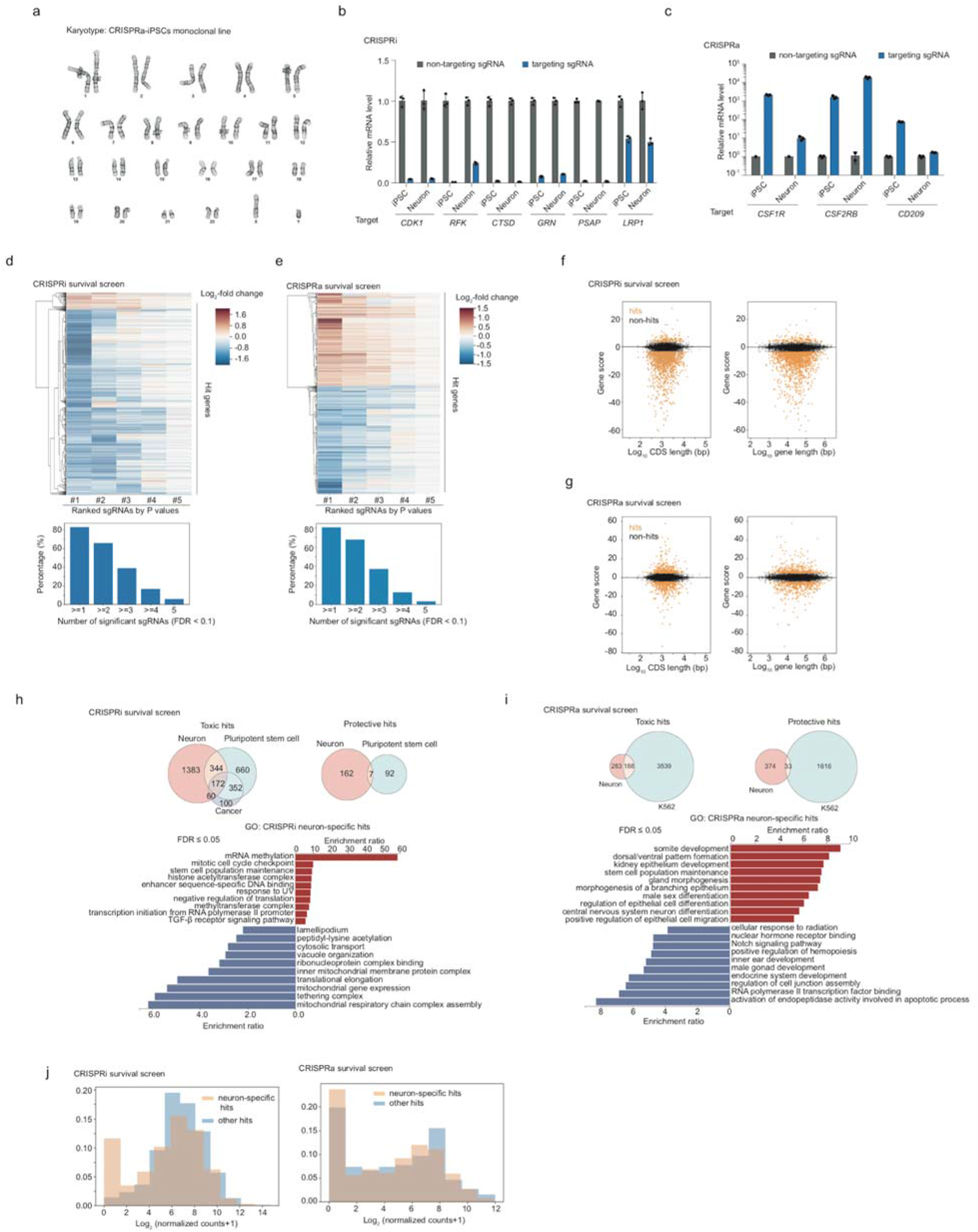
Karyotyping of the monoclonal CRISPRa-iPSC line, and analysis of CRISPRi and CRISPRa hits. (a) A normal karyotype was confirmed for the monoclonal CRISPRa-iPSC line. (b,c) Comparison of CRISPRi (b) and CRISPRa (c) efficacy in iPSCs and iPSC-derived neurons. The relative mRNA level of each targeted gene was calculated as the ratio of its expression in cells expressing a targeting sgRNA as compared to a non-targeting control sgRNA measured by qPCR (mean +/s sd, n = 3 technical replicates). The housekeeping gene ACTB was used for normalization. (d,e) Top, heatmaps showing phenotype scores (Log_2_-fold change) of all 5 sgRNAs (x-axis) targeting each hit gene (y-axis) from the primary CRISPRi (left) and CRISPRa (right) survival screens. The five sgRNAs targeting a given gene are ranked by the significance of their P values and are shown from left to right. Bottom, bar graphs summarizing the percentage of hit genes that have a certain number of sgRNAs (x-axis) showing a significant phenotype (false discovery rate (FDR) < 0.1) in CRISPRi (left) and CRISPRa (right) survival screens. (f,g) Scatter plots showing the relationship between Gene Score and gene coding sequence (CDS length (left) or gene length (right) for genome-wide CRISPRi (f) and CRISPRa (g) survival screens. (h, i) Top: Venn diagrams comparing CRISPRi (h) and CRISPRa (i) screen results for neuronal survival from this paper with other published survival screens for different human cell types. For CRISPRi, hit genes with toxic phenotypes for the survival of neurons were compared with those for cancer cells (‘gold-standard’ essential genes^14^) and pluripotent stem cells^11–13^ (genes that were identified as essential in more than one studies were retained for comparison). Protective hits for the survival of neurons were compared with those for human pluripotent stem cells^11,12^ (genes that were identified as essential in both studies were retained for comparison). For CRISPRa, hits were compared with our published survival screen in K562 cells^10^ reanalyzed using our MAGeCK-iNC pipeline. Bottom: Gene Ontology (GO) term enrichment analysis was conducted for hits resulting in increased survival (red) or decreased survival (blue); terms are shown up to an FDR of 0.05. (j) Neuronal expression levels of neuron-specific hit genes and other hit genes from CRISPRi (top) and CRISPRa (bottom) screens are shown, binned by order of magnitude.

**Extended Data Fig. 2:**
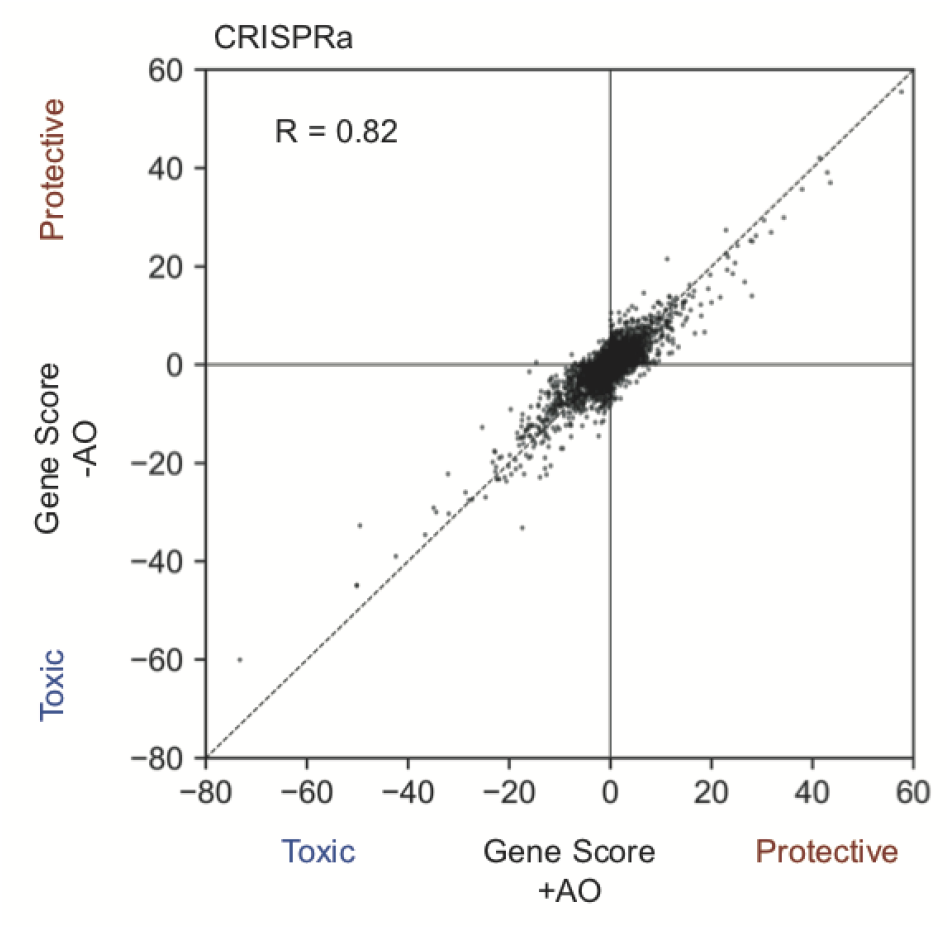
Comparing CRISPRa survival screens in +AO and -AO conditions. Each dot represents one gene, and its Gene Score in the +AO screen was plotted on the x-axis and Gene Score in the -AO screen on the y-axis. The Pearson correlation coefficient is shown.

**Extended Data Fig. 3:**
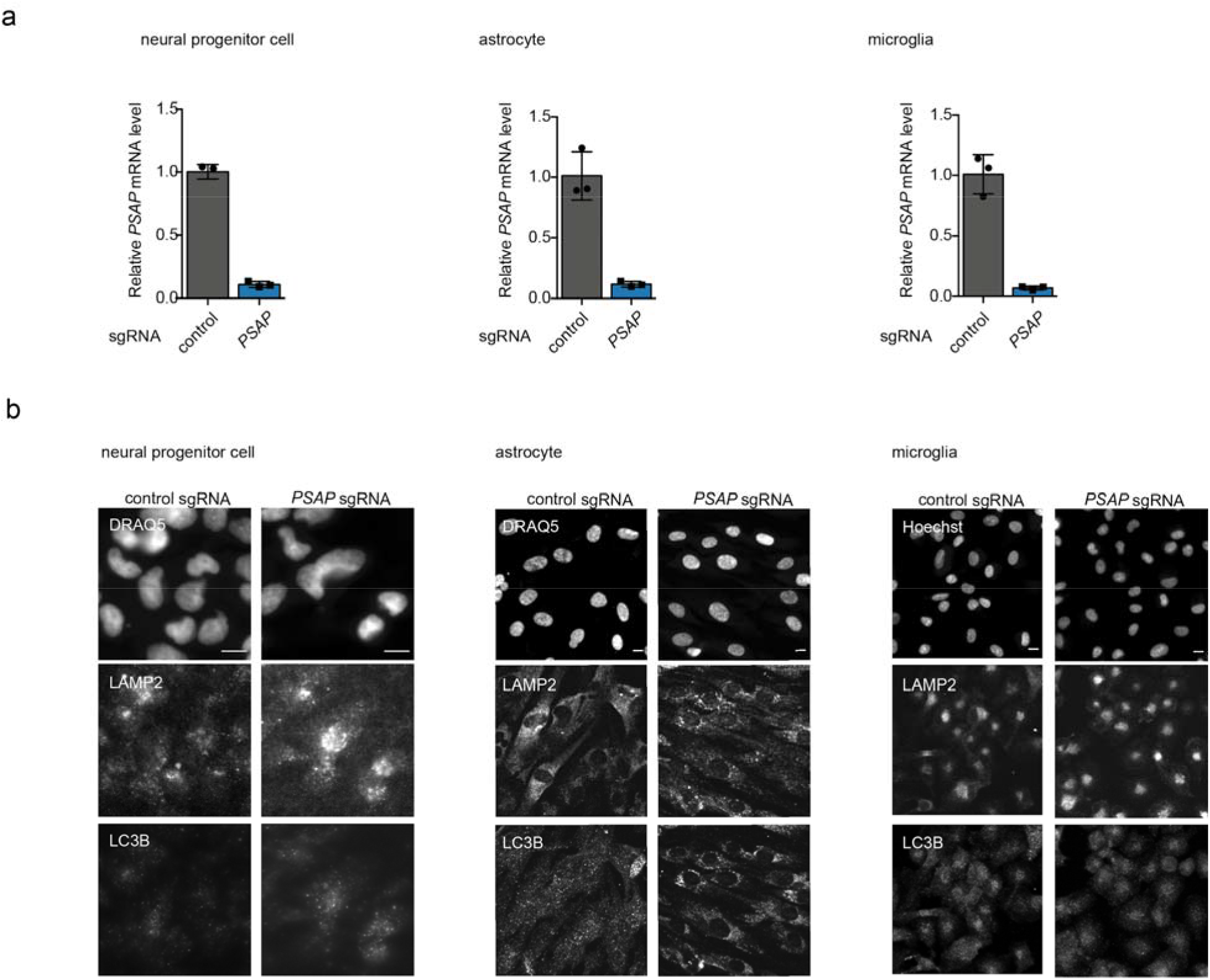
Characterization of *PSAP* KO in other cell types. (a) qPCR validation of *PSAP* knockdown in neural progenitor cells (left), astrocytes (middle) and microglia (right) diffentiated from CRISPRi iPSCs expression a *PSAP* sgRNA as compared to a non-targeting control sgRNA (mean +/s sd, n = 3 technical replicates). The housekeeping gene ACTB was used for normalization. (b) Representative fluorescence microscopy images for neural progenitor cells (left), astrocytes (middle) and microglia (right) diffentiated from CRISPRi iPSCs expression a non-targeting sgRNA or a *PSAP* sgRNA, stained with LAMP2 and LC3B antibodies. DRAQ5 was used for nuclear staining. Scale bar, 10 μm.

**Extended Data Fig. 4:**
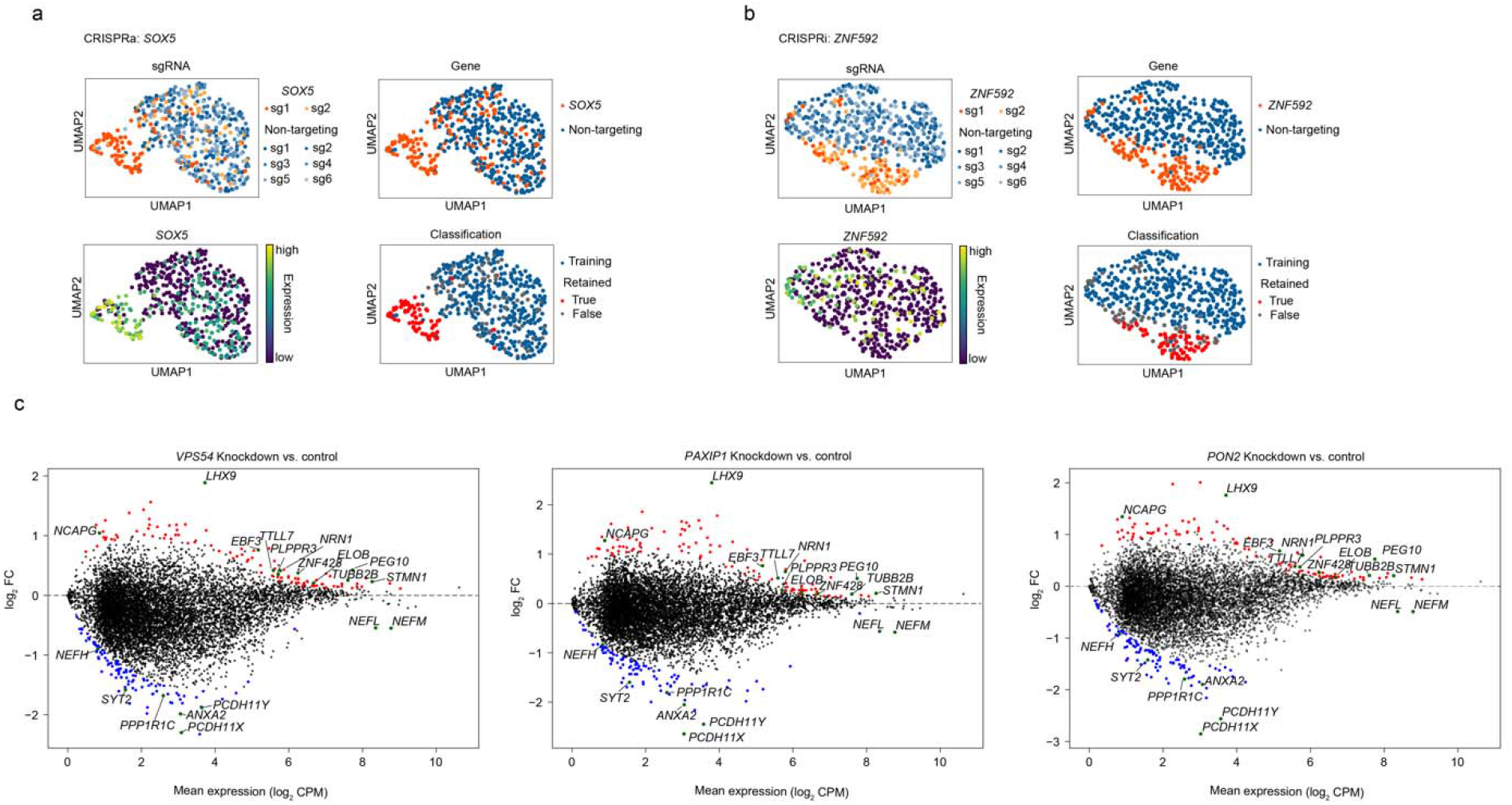
Examples of the CROP-seq classification method, and shared transcriptomic signatures of *VPS54, PAXIP1,* and *PON2* knockdown in human iPSC- derived neurons. (a,b) CROP-seq examples showing the application of the outlier detection-based classification method in cases where two sgRNAs targeting the same gene had heterogeneous efficacy (a, *SOX5* in CRISPRa) or the expression level of the target gene was too low to quantify knockdown level (b, *ZNF592* in CRISPRi). (c) Transcriptomic changes induced by knockdown of *VPS54* (left), *PAXIP1* (middle), and *PON2* (right) in neurons. For each perturbation, the top 200 upregulated and downregulated genes compared to control (i.e. unperturbed cells) are shown in red and blue, respectively. Within this set, shared genes among all three perturbations are highlighted in green.

**Supplementary Figure 1:**
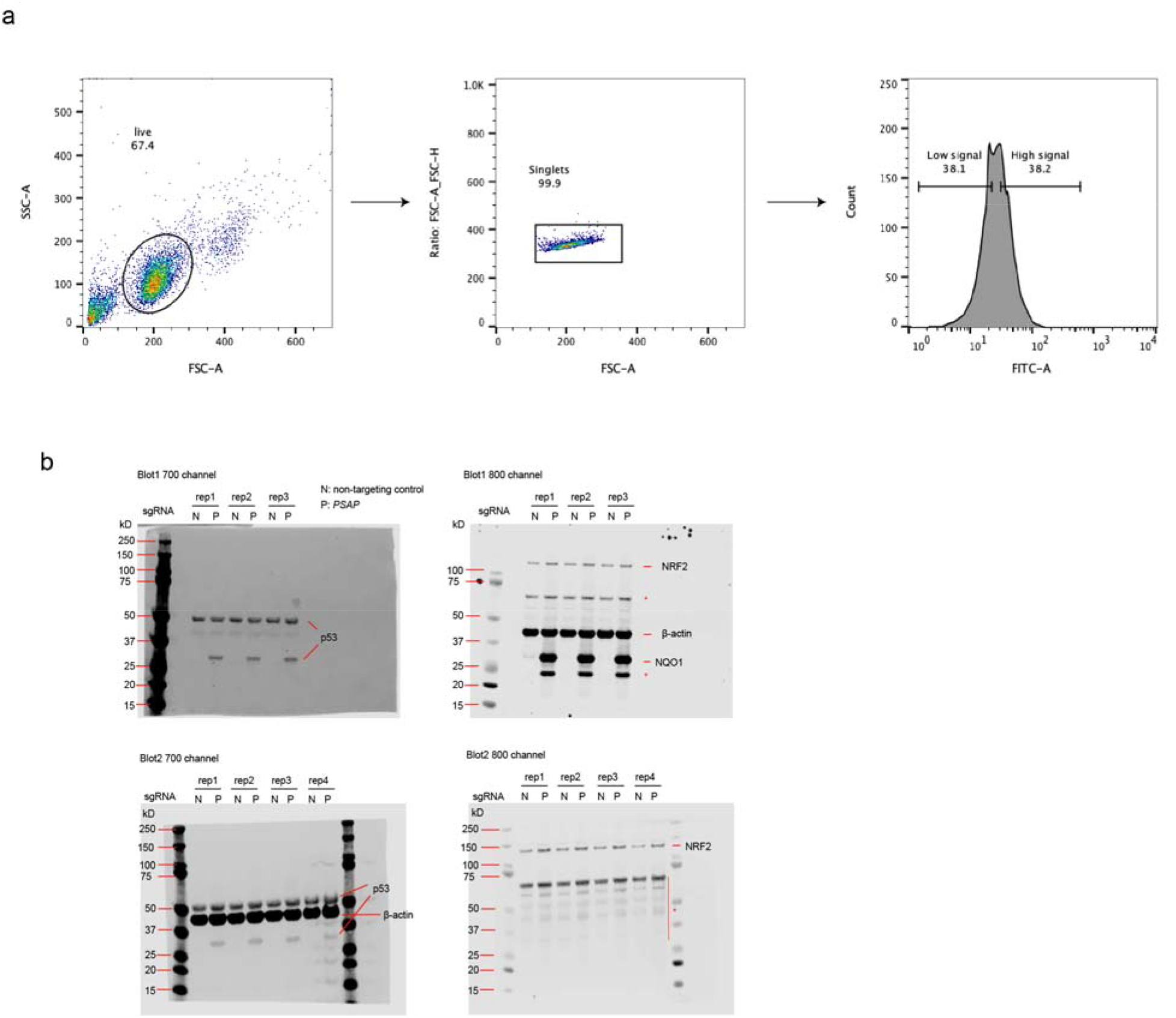
Gating strategy for FACS-based screens and original images of western blots. (a) Intact neurons were identified from the FSC-SSC plot and then gated for singlets. These cells were sorted into high and low signal populations corresponding to the top 40% and the bottom 40% of the staining signal distribution. (b) Original western blots images for Fig. 6e & f Lysates from CRISPRa neurons expressing the non-targeting control sgRNA or PSAP sgRNA were subjected to SDS-PAGE and Western blotting, and probed for p53, β-actin, NRF2 and NQO1 in two independent experiments. The blots were imaged using the LI-COR system in the 700 and 800 channels. Asterisks (*) label unspecific or indeterminate bands.

## Supplementary Table Legends

**Supplementary Table 1: Screen results for primary and pooled validation screens**

Screens were analyzed using the MAGeCK-iNC pipeline (see Methods for details). Hit class values of 1, −1 or 0 were assigned to hit genes with positive Phenotype scores, hit genes with negative Phenotype scores or non-hits, respectively. Each screen is provided in a separate tab.

**Supplementary Table 2: Hit class info for Fig2g**

This table shows the hit class info for genes in each screen in Fig2g. Hit class values of 1, −1 or 0 were assigned to hit genes with positive Phenotype scores, hit genes with negative Phenotype scores or non-hits, respectively.

**Supplementary Table 3: Untargeted lipidomics data for WT and *PSAP* KO neurons**

Untargeted lipidomics data for WT and *PSAP* KO neurons. P values were calculated using Student’s t-test and were corrected for multiple testing using the Benjamini-Hochberg method.

**Supplementary Table 4: sgRNA sequences for pooled validation and CROP-seq libraries** These tables show the protospacer sequences for sgRNAs in the pooled validation and CROP-seq libraries. Each library is provided in a separate tab. sgRNA information for the genome-wide libraries was previously published^10^.

**Supplementary Table 5: sgRNA cell counts for CROP-seq screens**

These tables summarize the number of cells for sgRNAs in the CRISPRi and CRISPRa CROP-seq screens. First tab: CRISPRi, second tab: CRISPRa.

## Notes

https://crisprbrain.org

### Summary of Updates

Additional experiments, analyses, and data tables. Two additional authors.

